# *De novo* design of therapeutic scFvs and multi-specific engagers from sequence alone

**DOI:** 10.64898/2026.03.17.712263

**Authors:** Takashi Fujiwara, Hideyuki Shimizu

**Affiliations:** Department of AI Systems Medicine, M&D Data Science Center, Institute of Integrated Research, Institute of Science Tokyo, Tokyo, JAPAN; Graduate School of Medical and Dental Sciences, Institute of Science Tokyo, Tokyo, JAPAN

**Keywords:** Generative antibody design, *De novo* biotherapeutic engineering, Structure-independent binder discovery, Computational developability assessment

## Abstract

The discovery of functional antibody therapeutics is fundamentally constrained by the astronomical complexity of the sequence landscape and the inefficiency of empirical screening. While structure-based computational design has made significant strides, its reliance on high-resolution co-complex data restricts its utility against the vast undruggable proteome and rapidly evolving pathogens. Here, we introduce IASO, a sequence-centric generative framework for the *de novo* design of diverse biotherapeutic modalities, including single-chain variable fragments and multi-specific engagers, requiring only antigen sequence as input. IASO integrates an evolutionary-informed generative engine that constructs antigen-steered candidate libraries with a high-fidelity interaction module capturing synergistic physicochemical motifs. We demonstrate the framework’s robustness by identifying novel binders against diverse clinical targets, where IASO successfully discriminates atomic-level mutations to overcome established drug resistance in EGFR and distinguishes single-residue variants of SARS-CoV-2. Furthermore, we extend this approach to the *in silico* construction of a complex bispecific T-cell engager that exhibits developability profiles comparable to clinically approved therapeutics, including enhanced solubility and minimal immunogenicity. By transforming serendipitous exploration into predictable high-resolution engineering, IASO establishes a scalable foundation for the rapid development of next-generation biopharmaceuticals directly from sequence data alone.

## Introduction

Monoclonal antibodies and their derivative formats have emerged as dominant therapeutic modalities with over 150 approved drugs generating annual revenues exceeding $200 billion and accelerating clinical applications across oncology, immunology, and infectious diseases^1–3^. The rapid expansion of this therapeutic landscape has driven intensive research into next-generation modalities, particularly single-chain variable fragments (scFvs)^4^. Despite their compact size, they retain high antigen specificity and broad applicability while offering decisive advantages including chemical synthesizability without relying on complex cell culture systems and superior tissue penetration^5,6^. These properties position scFvs as critical building blocks for advanced therapeutics such as bispecific T-cell engagers (BiTEs)^7–9^ and chimeric antigen receptor (CAR) T-cell therapies^10–12^. Consequently, advances in scFv discovery have implications that extend beyond single binders to the broader engineering of multi-specific and cell-engaging therapeutics. However, the discovery of functional scFvs remains a formidable challenge because even this compact antibody format occupies an enormous combinatorial sequence space^13^, whereas empirical screening and selection processes are inherently stochastic and susceptible to sampling bias^14^. Traditional platforms including hybridoma technology, phage display, and yeast surface display typically require labor-intensive, iterative screening and optimization^15^, creating critical bottlenecks in responding to emerging pathogens and evolving drug resistance, thereby highlighting an urgent need to streamline the design process^16,17^.

To address these bottlenecks, deep learning and protein language models (PLMs) are increasingly being leveraged to advance computational scFv discovery, enabling both accurate prediction of antibody–antigen interactions (AAI) and the *de novo* design of functional binding sequences^18–20^. Foundation models such as ESM-2^21^ and specialized antibody language models, including AbLang-2^22^, have substantially improved sequence representation learning, and recent predictors such as RLEAAI^23^ and DeepInterAware^24^ have further enhanced AAI prediction by integrating latent embeddings with attention-based neural architectures^25^. However, progress remains constrained by the scarcity of high-quality AAI data and pervasive biases toward well-characterized model antigens, both of which compromise generalization to real-world screening settings^26^. Furthermore, many existing predictors remain only loosely connected to generative design and downstream validation workflows, limiting their utility in end-to-end scFv discovery pipelines^18^.

Regarding *de novo* design, recent advances in generative modeling have progressively shifted the field from antigen-agnostic antibody sequence generation toward target-specific design. Large-scale antibody generative models such as ProGen2-OAS^27^ can generate diverse and natural-like paired antibody sequences, but generally lack explicit antigen conditioning, limiting their utility for target-directed scFv discovery. More recently, sequence-based target-specific approaches such as MAGE^28^ have demonstrated that scFv generation can be conditioned directly on antigen information without requiring an experimentally resolved antigen–antibody complex structure. In parallel, structure-aware methods such as BoltzGen^29^ and RFantibody^30,31^ have highlighted the potential of antigen-structure-informed binder design, but their applicability is inherently constrained by the need for reliable structural information and high-quality template scaffolds to support effective design^32^. Together, these developments represent important steps toward computational scFv discovery. However, existing approaches remain fragmented, with target-conditioned generation, interaction prediction, and downstream structural or pharmaceutical evaluation rarely integrated within a unified end-to-end framework^33–35^. Consequently, current methods provide only limited support for the reliable generation and prioritization of functional scFvs, and the identification of candidates with favorable binding potential, structural stability, and pharmaceutical properties still depends heavily on downstream screening and experimental triage^36–38^.

Here we present IASO (*In silico* Antigen-conditioned scFv Optimization), an integrated end-to-end framework that addresses these limitations through the synergistic combination of three computational modules operating exclusively on primary sequence data. IASO comprises IASO-Gen, an evolutionary-informed generative engine that produces target-specific antibody libraries through prefix-tuning and low-rank adaptation of protein language models, IASO-AAI, a high-precision interaction predictor that captures synergistic physicochemical motifs by integrating antibody-specific evolutionary embeddings with local compositional features, and a structure-based validation module that uses state-of-the-art (SOTA) complex prediction models for final candidate refinement. We show that IASO-AAI substantially outperforms existing AAI prediction models across multiple benchmarks while maintaining exceptional calibration, and that IASO-Gen generates diverse and developable candidates with favorable physicochemical properties, immunogenicity profiles, and structural features that closely resemble those of natural human antibody repertoires. Through comprehensive case studies focused on clinically critical challenges, including drug-resistant EGFR mutants that abolish binding of approved therapeutics and fine-grained SARS-CoV-2 variants differing by single residues, we show that the integrated pipeline achieves atomic-level design precision that has previously been accessible only through structure-guided approaches or extensive experimental screening. Beyond standalone scFv discovery, IASO is readily extensible to multi-specific formats constructed from scFvs. Successful application to BiTE design further demonstrates its versatility across therapeutic modalities, with designed molecules exhibiting developability profiles exceeding those of clinically approved BiTE therapeutics. By transforming stochastic exploration into predictable and high-resolution computational engineering, IASO establishes a new paradigm for rapid antibody discovery that is particularly well suited to urgent medical needs, including pandemic preparedness and the treatment of evolving drug resistance in precision oncology.

## Results

### IASO-AAI enables high-precision binder prioritization within the IASO pipeline

To address the fragmentation of existing computational scFv discovery strategies, we developed IASO, an integrated end-to-end framework for sequence-based scFv generation, prioritization, and refinement. IASO begins from target antigen sequences, from which IASO-Gen generates target-conditioned scFv libraries, IASO-AAI prioritizes candidate binders through antibody–antigen interaction (AAI) prediction, and a downstream structural filtering module refines the top-ranked molecules to yield final scFv candidates. This modular design enables the seamless coupling of generation, interaction-based ranking, and structural validation within a unified discovery workflow (**Fig. 1a**).

**Figure 1.**
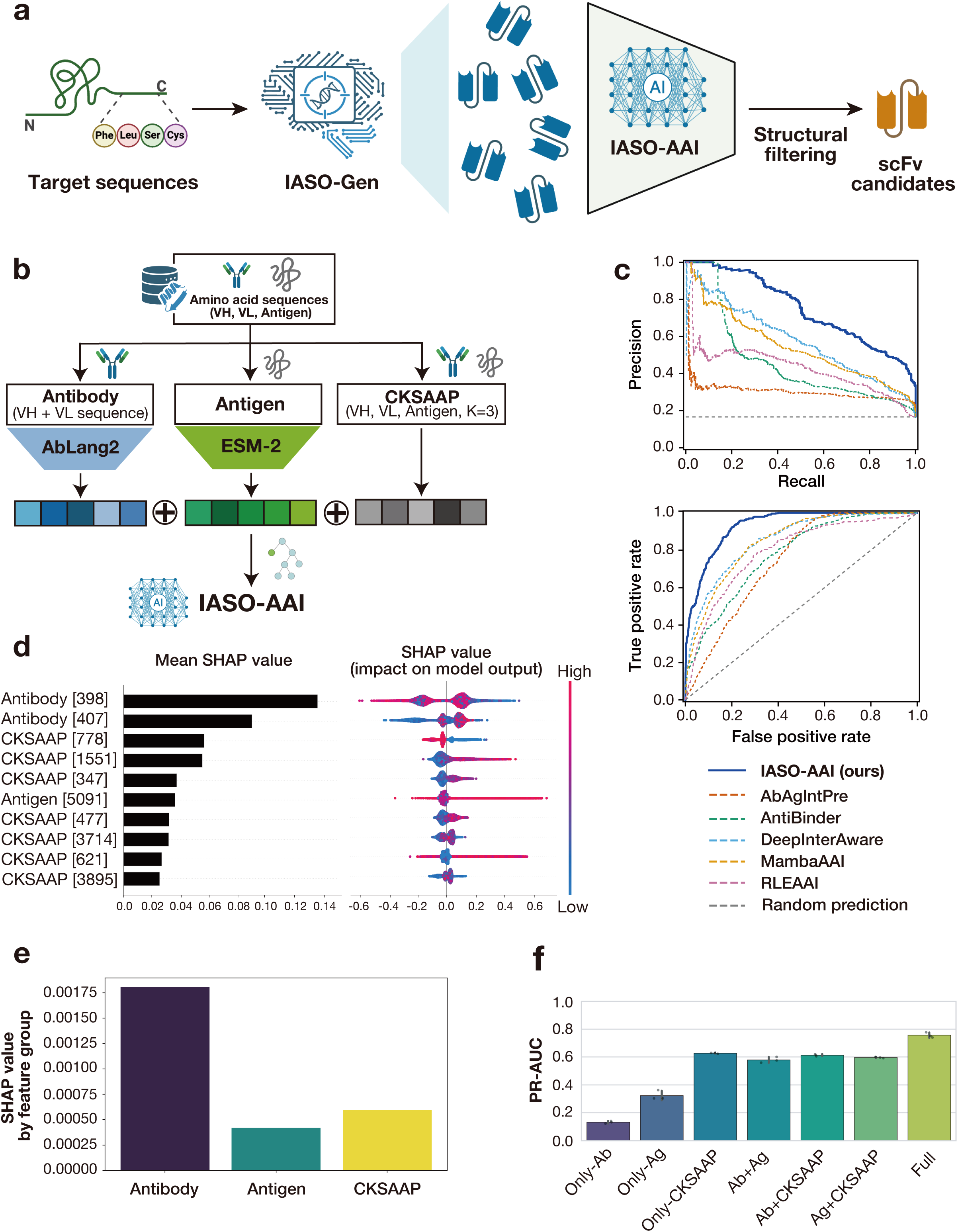
Architecture and performance of IASO-AAI. **(a)** IASO discovery workflow. Antigen sequence input undergoes sequential processing: IASO-Gen generates candidate antibodies, IASO-AAI predicts binding, and Boltz-2 evaluates structural compatibility. **(b)** IASO-AAI architecture. Antibody sequences (VH, VL) are embedded via AbLang-2 (480 dimensions), antigens via ESM-2 (5,120 dimensions), and both integrated with CKSAAP physicochemical features (4,800 dimensions) to predict binding probability. **(c)** Performance comparison against existing methods (AbAgIntPre, AntiBinder, DeepInterAware, MambaAAI, and RLEAAI). Top: Precision-Recall curves; bottom: Receiver Operating Characteristic (ROC) curves. IASO-AAI (blue) achieves superior performance on both metrics. **(d)** Feature importance via SHAP analysis. Bar and beeswarm plots show the top 10 individual features contributing to predictions. Bracketed numbers indicate dimension index within each feature set. **(e)** Mean absolute SHAP values across feature groups (antibody, antigen, and CKSAAP) reveal antibody embeddings contribute most significantly to interaction predictions. **(f)** Ablation study. Precision-Recall Area Under the Curve (PR-AUC) for models trained with different feature combinations (Ab: AbLang-2, Ag: ESM-2, CKSAAP) demonstrates that integrating all three feature types maximizes predictive performance. Error bars represent standard deviation across three independent runs with different random seeds.

Within this pipeline, efficient identification of true binders from vast sequence space represents a major bottleneck in therapeutic development. Therefore, we first developed IASO-AAI, a machine-learning framework for rapid, high-precision prediction of antibody–antigen interactions that serves as the binder-prioritization engine within IASO. IASO-AAI takes paired VH and VL sequences together with the target antigen sequence as input, allowing binding prediction to be performed in the context of the VH–VL pair rather than for each chain separately. As scFvs preserve the same paired variable-domain architecture in a single-chain format, IASO-AAI is directly applicable to scFv–antigen interaction prediction.

To support robust AAI prediction with high-quality, minimally biased training data, we curated a comprehensive dataset comprising 26,040 antibody–antigen interaction pairs from the AACDB^39^ and AbRank^40^ databases. To enable rigorous evaluation of generalization, antigen redundancy was removed using CD-HIT^41^, thereby reducing sequence-overlap bias across data partitions. In addition, following established computational benchmarks^42,43^, we adopted a 1:5 positive-to-negative ratio to emulate realistic screening scenarios in which true binders are rare relative to the large background of non-binding candidates (**Supplementary Fig. 1a**).

IASO-AAI was designed to integrate complementary sequence representations that capture both evolutionary constraints and physicochemical determinants of molecular recognition (**Fig. 1b**). We reasoned that antibody-specific language models would better represent the sequence grammar of immunoglobulins than general protein language models, and therefore encoded paired antibody sequences using AbLang-2^22^, while antigen sequences were represented using ESM-2^21^. To complement these global representations with features capturing local interaction determinants, we introduced Composition of k-Spaced Amino Acid Pairs (CKSAAP) features^44^, which quantify amino acid pair frequencies and capture local sequence patterns associated with physicochemical interaction determinants such as hydrophobic and electrostatic complementarity^45–47^. Parameter sensitivity analysis confirmed robust performance across *k* values (**Supplementary Fig. 1b**), with *k* = 3 selected following established benchmarks. This multi-scale integration was designed to link macro-level evolutionary patterns with micro-level physicochemical features that shape molecular recognition. Hyperparameter optimization via Bayesian search maximized predictive performance (**Supplementary Table 1**).

Performance evaluation on the independent test dataset demonstrated that IASO-AAI significantly outperformed five established methods including SOTA approaches^23,24,46,48,49^, achieving the highest performance among all compared models with a PR-AUC of 0.75, ROC-AUC of 0.93, and Precision of 0.8525 (**Fig. 1c, Supplementary Fig. 1c, Supplementary Table 2**). Notably, calibration analysis demonstrated excellent reliability, with a Brier score^50^ of 0.079 and an expected calibration error (ECE)^51,52^ of 0.049, indicating that model outputs represent accurate binding probabilities rather than arbitrary confidence scores (**Supplementary Fig. 1d**). This calibration is critical for practical applications, as it enables researchers to set meaningful thresholds for candidate selection. To verify that superior performance stemmed from the model architecture rather than dataset-specific overfitting, we performed external validation on an independent RLEAAI benchmark^23^. IASO-AAI achieved extremely high benchmarks (PR-AUC: 0.974, ROC-AUC: 0.969, F1 score: 0.907), surpassing all compared methods^23,43,46,53,54^ (**Supplementary Table 3**), confirming robust generalization across datasets.

To elucidate the rationale behind the predictive superiority of IASO-AAI, we conducted feature importance analysis using Shapley Additive Explanations (SHAP)^55^. Individual feature-level analysis revealed that the top-ranked features predominantly originated from the antibody embedding space, suggesting that immunoglobulin evolutionary patterns encode critical binding information (**Fig. 1d**). Aggregating SHAP values across feature categories confirmed this pattern, with antibody-specific evolutionary embeddings contributing most strongly to predictions. Mean SHAP values for antibody embeddings exceeded other feature groups by over threefold (**Fig. 1e**). Ablation studies confirmed that all three feature types are necessary for optimal performance, with models lacking any component showing substantially reduced performance (**Fig. 1f**). The synergistic contribution of evolutionary embeddings and physicochemical characteristics demonstrates that IASO-AAI achieves high precision by integrating information across multiple scales, transcending simple sequence pattern recognition to capture the biophysical principles governing antibody-antigen recognition.

### IASO-Gen generates diverse, antigen-conditioned scFv sequences

While IASO-AAI provides high-precision discrimination of binders, accelerating scFv discovery requires autonomous generation of candidate sequences^28,33^. Therefore, we developed IASO-Gen, a generative framework that designs scFv sequences conditioned solely on target antigen sequences. The central challenge was balancing preservation of immunoglobulin structural grammar with target-specific binding requirements. To address this, we built upon ProGen2-OAS^27^, which encodes antibody sequence rules through large-scale fine-tuning of ProGen2 on the Observed Antibody Space (OAS) database^56^, but lacks antigen conditioning capability. We introduced Prefix-tuning^57,58^ to inject antigen information into the Transformer architecture and employed Low-Rank Adaptation (LoRA)^59^ to enhance sequence expressivity with limited data sources (**Fig. 2a, see Materials and Methods**).

**Figure 2.**
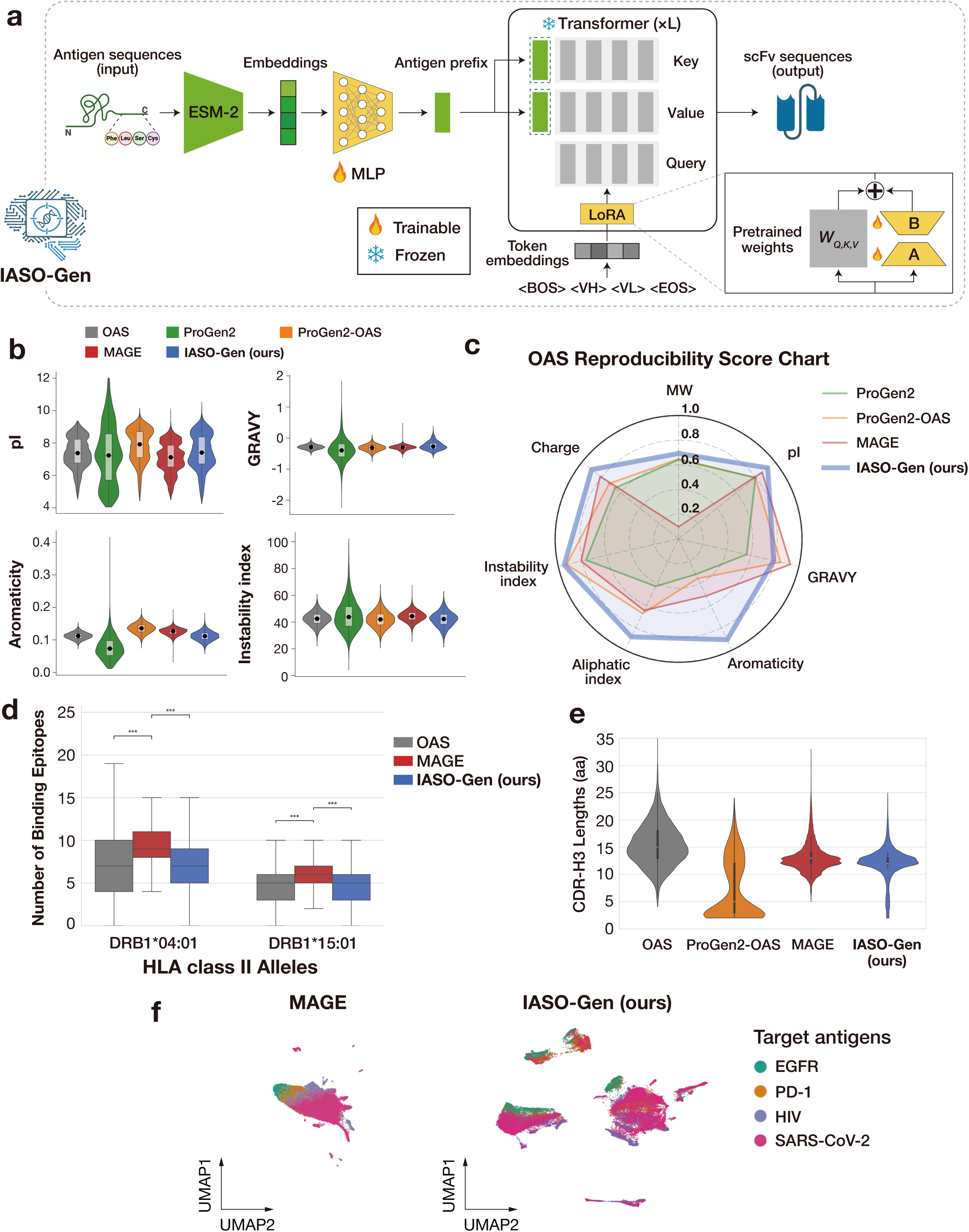
IASO-Gen architecture and characterization of generated antibodies. **(a)** IASO-Gen architecture. Antigen sequences are encoded by ESM-2 and projected by an MLP into antigen prefixes, which are injected into the Key and Value streams of the ProGen2-OAS transformer blocks. LoRA modules applied to the transformer attention layers enable parameter-efficient fine-tuning for antigen-conditioned scFv generation. **(b)** Physicochemical properties for 10,000 antibodies from the OAS database (reference) and four generative models (ProGen2, ProGen2-OAS, MAGE, and IASO-Gen). Violin plots compare molecular weight, isoelectric point (pI), hydrophobicity (GRAVY), aromaticity, aliphatic index, instability index, and net charge. **(c)** OAS Reproducibility Score based on the Kolmogorov-Smirnov statistic for each physicochemical property. IASO-Gen (blue) closely matches the natural antibody distribution. **(d)** Immunogenicity assessment. Predicted strong HLA-II binders for two HLA-DRB1 alleles (DRB1*04:01, DRB1*15:01). IASO-Gen (blue) shows low epitope burden comparable to OAS (gray), whereas MAGE (red) exhibits significantly elevated risk (\*\*\**p* < 0.001). **(e)** CDRH3 length distributions. IASO-Gen reproduces the natural length profile from OAS (gray), while ProGen2-OAS (orange) generates shorter loops. **(f)** UMAP projection of antibody sequence embeddings generated under four antigen conditions (EGFR, PD-1, HIV, SARS-CoV-2). MAGE (left) produces largely overlapping distributions, whereas IASO-Gen (right) generates distinct, antigen-specific clusters.

We next assessed whether the generated sequences exhibited physicochemical properties and safety-related characteristics comparable to those of natural human antibodies, supporting their developability as pharmaceutical candidates. Analysis of seven properties (molecular weight, isoelectric point (pI), GRAVY hydrophobicity^60^, aromaticity, aliphatic index^61^, instability, net charge) revealed that IASO-Gen maintains high fidelity to the natural OAS repertoire^56^ (**Fig. 2b, Supplementary Fig. 2a**). Quantifying distributional similarity via the OAS Reproducibility Score (defined as 1 minus the Kolmogorov-Smirnov (KS) statistic, see **Materials and Methods**), IASO-Gen outperformed existing models including MAGE^28^, confirming generation of biologically realistic sequences (**Fig. 2c**).

Beyond physicochemical properties, minimizing immunogenicity risk is critical for clinical translation^62^. Human Leukocyte Antigen (HLA) class II binding predictions via NetMHCIIpan 4.3^63^ showed that IASO-Gen sequences exhibit low predicted epitope burden comparable to natural antibodies, whereas MAGE^28^ showed significantly elevated risk (**Fig. 2d**). For this analysis, we prioritized HLA-DRB1*04:01 and HLA-DRB1*15:01, as these HLA class II alleles are highly clinically relevant and have been extensively associated with human immune response and disease susceptibility^64–66^. IASO-Gen maintained low predicted epitope burden comparable to natural antibodies across these allelic contexts, whereas MAGE^28^ exhibited significantly elevated risk. This reduced immunogenicity likely reflects IASO-Gen’s closer adherence to natural sequence distributions.

Structural validity was further assessed through CDRH3 analysis, as this hypervariable region determines binding specificity^67^ yet must conform to physiological length constraints for conformational stability^68^. IASO-Gen accurately reproduced natural CDRH3 length distributions (**Fig. 2e**), confirming structurally sound design. Sequence diversity analysis demonstrated that IASO-Gen maintains higher positional Shannon entropy^69^ in CDR loop centers compared to existing models (**Supplementary Fig. 2b**) and shows larger pairwise Levenshtein distances^70,71^ than MAGE^28^ (**Supplementary Fig. 2c**), indicating broad sequence space exploration without mode collapse. These results establish that IASO-Gen balances structural realism with the diversity needed for therapeutic candidate identification.

The critical test of IASO-Gen is whether it generates target-specific sequences rather than generic antibodies. We performed UMAP projection^72^ of antibody sequence embeddings generated under different antigen conditions. While unconditioned ProGen2^27^ and MAGE^28^ produced largely overlapping clusters, IASO-Gen formed distinct, well-separated clusters when conditioned on EGFR, PD-1, HIV gp120, or SARS-CoV-2 spike (**Fig. 2f, Supplementary Fig. 2d**). This antigen-specific clustering demonstrates that IASO-Gen captures fine-grained antigen features to enable tailored library design for each target.

Together, these validations establish that IASO-Gen possesses essential attributes for therapeutic development, combining biological safety, sequence diversity, and target specificity to enable rational, antigen-directed scFv design.

### Integrated IASO pipeline enables design of EGFR resistance-overcoming scFvs

Having shown that IASO-AAI enables high-precision binder prioritization (**Fig. 1)** and that IASO-Gen generates diverse, antigen-conditioned scFv libraries with favorable developability-related properties (**Fig. 2**), we next asked whether combining these two modules within the IASO workflow (**Fig. 1a**) could enable practical therapeutic scFv discovery in a realistic design setting. To address this, we applied the full IASO workflow to EGFR, executing sequential antigen-conditioned generation, interaction-based screening, and structure-based filtering (**Fig. 3a**). This staged strategy allows large candidate libraries to be progressively narrowed to a small set of high-confidence binders, thereby coupling scalability with structural rigor while minimizing computational cost. To demonstrate practical utility, we targeted the Epidermal Growth Factor Receptor (EGFR), a critical oncology target with well-characterized drug-resistant mutations including S468R^73^. This mutation abolishes binding of the clinical antibody Cetuximab^74^, presenting a therapeutically relevant challenge for computational design. We generated 20,000 EGFR-targeting scFv sequences with IASO-Gen and subsequently filtered them by successful ANARCI-based annotation, retaining 10,990 sequences with valid antibody numbering for downstream analysis. Screening of this quality-controlled library with IASO-AAI identified 116 predicted binders (**Fig. 3b**). Notably, IASO-Gen-derived sequences showed significantly higher IASO-AAI binding scores compared to baseline models (**Fig. 3c**), confirming that antigen conditioning enriches for high-affinity candidates. Structural evaluation of top-ranked sequences using Boltz-2^75^ revealed superior structural quality, as indicated by higher confidence scores (pLDDT, pTM) and lower prediction error (pde), compared to both baseline-generated sequences (**Fig. 3d**) and a previously reported anti-EGFR scFv^76^ (**Fig. 3e**), validating the pipeline’s multi-stage filtering strategy.

**Figure 3.**
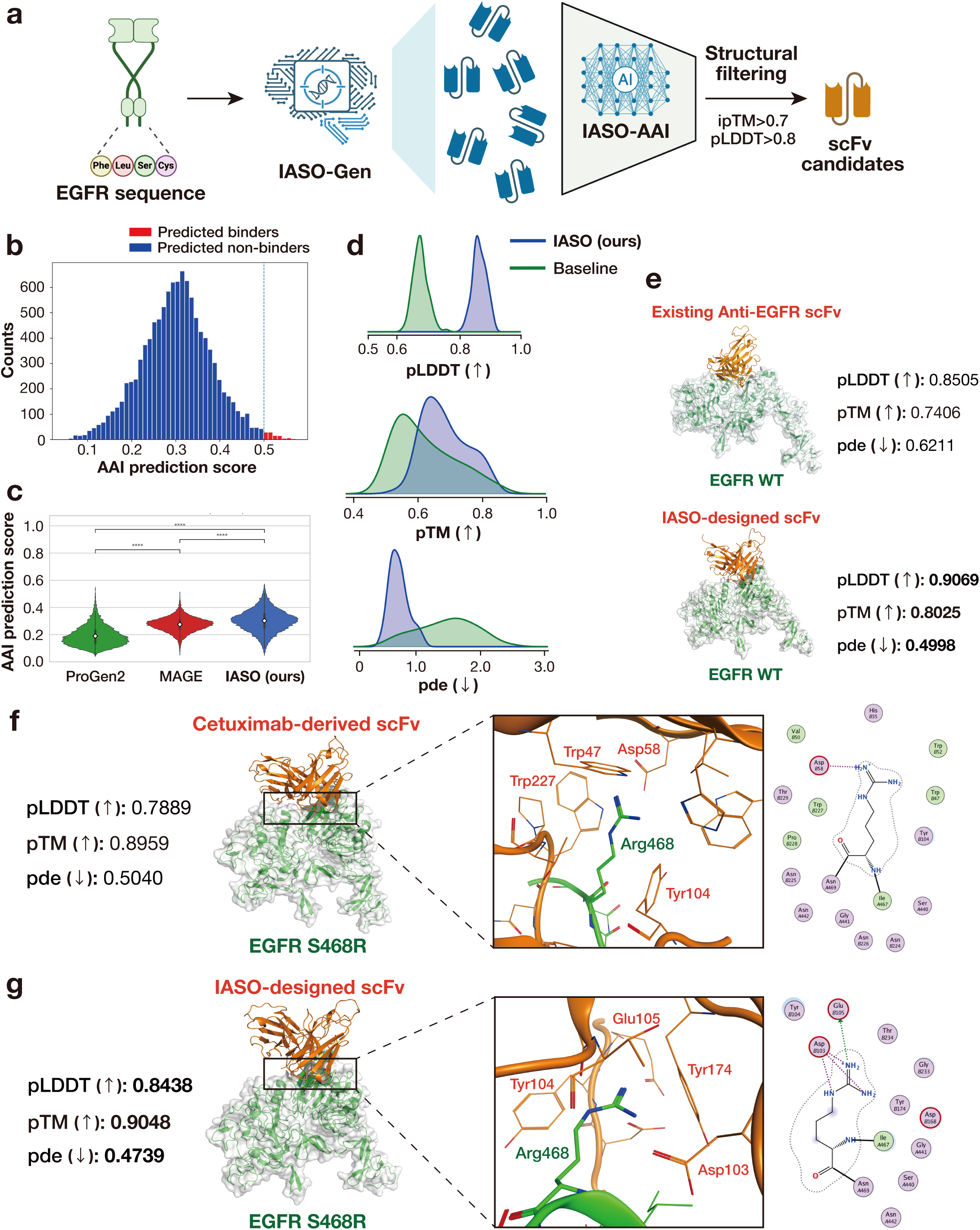
EGFR-targeting antibody design via the integrated IASO pipeline. **(a)** Schematic overview of the integrated IASO workflow for EGFR-targeting scFv design. Starting from the EGFR sequence, IASO-Gen generates an antigen-conditioned scFv library, IASO-AAI prioritizes candidate binders by antibody– antigen interaction prediction, and downstream structural filtration refines the library to yield high-confidence EGFR-targeting scFv candidates. **(b)** Binding score distribution for the IASO-Gen-generated anti-EGFR library. Sequences are classified as binders (red) or non-binders (blue) using a 0.5 threshold. **(c)** Comparative binding score distributions for scFvs from IASO-Gen, MAGE and ProGen2. IASO-Gen produces higher-scoring candidates (**** *p* < 0.0001). **(d)** Structural confidence metrics (pLDDT, pTM, pde) from Boltz-2 for the top 100 candidates. IASO-Gen candidates (blue) exhibit superior structural quality (higher pLDDT and pTM, lower pde) compared to baseline model (ProGen2, green). **(e)** Structural comparison of Boltz-2-predicted complexes. A previously reported anti-EGFR scFv^76^ (top) versus IASO-designed lead candidate (bottom), showing improved structural metrics (higher pLDDT/pTM, lower pde) for the IASO design. **(f)** Binding mode analysis for the EGFR S468R resistance mutant. Cetuximab-derived scFv shows steric clashes between Arg468 (green) and paratope residues Trp47/Trp227/Tyr104, compromising binding. **(g)** IASO-designed scFv forms optimized electrostatic interactions via Asp103 and Glu105 that accommodate Arg468. Left to right: Boltz-2 confidence metrics (pLDDT, pTM, pde), overall complex structure, detailed view of Arg468 environment (red/blue spheres: oxygen/nitrogen), and 2D interaction map (red: acidic, blue: basic, pink: polar, green: hydrophobic residues, purple line: ionic contact, green arrow: sidechain donor/acceptor).

We next challenged the pipeline to design scFvs targeting the S468R resistance mutant. Structural prediction revealed that IASO-designed candidates achieve higher structural confidence than Cetuximab-derived scFv when bound to the mutant (**Figs. 3f and 3g)**. Atomic-level analysis elucidated the mechanistic basis for this improved compatibility. In Cetuximab-derived scFv, the S468R mutation introduces severe steric clashes between Arg468 and bulky paratope residues Trp47, Trp227, and Tyr104, disrupting binding (**Fig. 3f**). In contrast, the IASO-designed scFv accommodates Arg468 through optimized electrostatic interactions with Asp103 and Glu105, establishing a binding mode compatible with the mutant epitope (**Fig. 3g, Supplementary Fig. 3a**). These results suggest that the IASO pipeline captures subtle structural perturbations to generate resistance-overcoming leads through rational design.

Furthermore, we evaluated the developability and safety profiles using Cetuximab-derived scFv as a clinical benchmark. Aggregation propensity analysis (Aggrescan4D)^77^ indicated that the IASO-designed scFv (**Fig. 3g**) maintains favorable scores comparable to the approved drug (**Supplementary Fig. 3b**). Solubility predictions via the CamSol method^78^ exceeded the threshold for pharmaceutical viability (**Supplementary Fig. 3c**), and immunogenicity assessment via NetMHCIIpan 4.3^63^ revealed HLA class II binding frequencies at or below Cetuximab-derived scFv levels across major HLA class II alleles^63,79,80^ (**Supplementary Fig. 3d**). For immunogenicity assessment, we focused on a representative panel of 12 major HLA-DRB1 alleles, because HLA-DRB1 has historically been the primary HLA class II locus used in immunogenicity evaluation^81^. Together, these multifaceted results strongly support the IASO as a robust platform for autonomous design of biopharmaceutical-grade scFv development.

### IASO enables versatile scFv design across diverse antigens and viral variants

To evaluate generalizability beyond oncology targets, we applied IASO to structurally diverse antigens spanning cancer immunotherapy (PD-1)^82^, infectious disease (HIV gp120, SARS-CoV-2 receptor-binding domain (RBD) variants), and host receptors (ACE2). Across all targets, IASO-designed candidates showed consistently favorable scFv–antigen complex prediction metrics, supporting the robustness of the framework across diverse antigen classes (**Fig. 4a, Supplementary Fig. 4a**). Together, these results indicate that IASO operates according to antigen-agnostic design principles and is broadly applicable as a versatile framework for scFv discovery.

**Figure 4.**
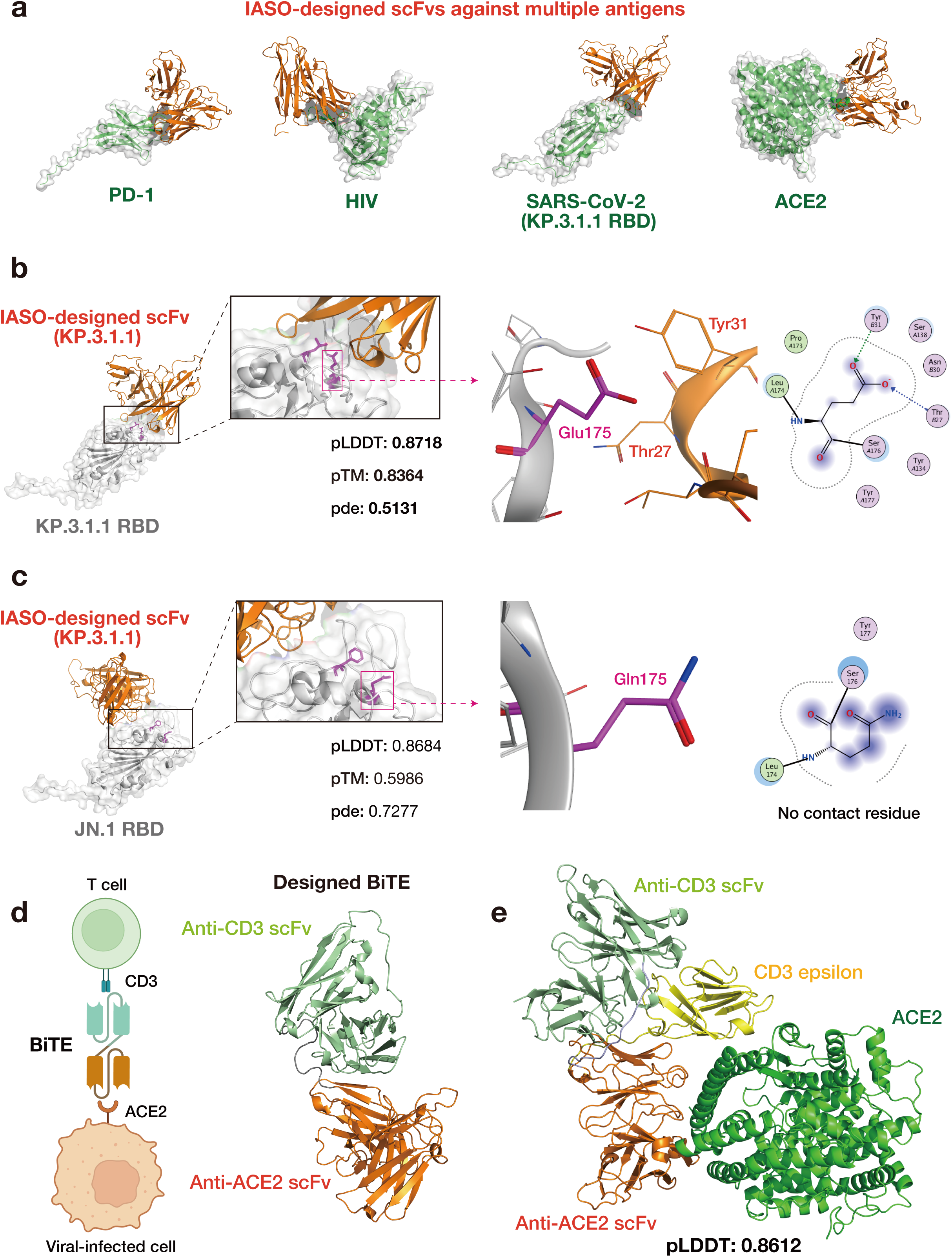
IASO versatility across diverse antigens and multi-specific formats. **(a)** Predicted complexes of IASO-designed scFvs (orange) with clinically relevant antigens (green): PD-1, HIV gp120, SARS-CoV-2 KP.3.1.1 RBD, and ACE2. Target antigens are shown as cartoon representations overlaid with semi-transparent molecular surfaces (gray). **(b)** Variant discrimination for SARS-CoV-2. Left: scFv designed against the KP.3.1.1 RBD (orange) bound to KP.3.1.1 (gray), showing high structural confidence (pLDDT 0.8718, pTM 0.8364; higher is better). Right: same scFv with ancestral JN.1 strain (gray), showing substantially reduced confidence (pTM 0.5986), demonstrating variant-specific selectivity. Mutation site in magenta. Close-ups show the binding interface. **(c)** Atomic-level interactions at residue 175 (magenta). Left: Glu175 (KP.3.1.1) forms hydrogen bonds with scFv Tyr31 (orange). Right: Gln175 (JN.1) lacks stabilizing contacts. Red/blue spheres: oxygen/nitrogen atoms. 2D interaction maps: red (acidic), blue (basic), pink (polar), green (hydrophobic), blue arrow (backbone donor/acceptor), green arrow (sidechain donor/acceptor), blue blurred outline (ligand exposure). **(d)** BiTE architecture. Anti-CD3 scFv (light green, T-cell recruitment) linked to IASO-designed anti-ACE2 scFv (orange, target recognition) via a flexible linker. **(e)** Ternary complex structure. The BiTE simultaneously engages the CD3 epsilon chain (yellow) and ACE2 (green) with high structural confidence (pLDDT 0.8612).

This breadth enables rapid response to viral immune escape mutations. We tested whether IASO could discriminate between closely related SARS-CoV-2 Omicron sublineages JN.1^83^ and KP.3.1.1^84^, which differ by only two RBD residues yet exhibit distinct evolutionary trajectories. ScFvs designed to target the KP.3.1.1 RBD maintained high structural confidence when modeled with the intended target but showed substantially reduced confidence scores with ancestral JN.1, despite preserved local folding (**Fig. 4b**). This specificity stems from atomic-level differences at position 175, where Glu175 (KP.3.1.1) forms favorable hydrogen bonds with the IASO-designed scFv, while Gln175 (JN.1) fails to establish stabilizing contacts (**Figs. 4b and 4c**). These results demonstrate that IASO achieves single-residue discrimination, enabling immediate design of variant-specific scFvs upon emergence of new strains.

### IASO pipeline extends to multi-specific scFvs engineering

Beyond single-chain formats, we explored IASO’s capacity to design Bispecific T-cell Engagers (BiTEs), which require coordinated function of multiple scFv domains. While IASO can rapidly generate variant-specific scFvs, targeting the evolutionarily conserved host receptor ACE2 offers a mutation-resistant therapeutic strategy against rapidly evolving viruses such as SARS-CoV-2^85,86^. We therefore designed a BiTE to redirect T cells to ACE2-expressing infected cells for targeted cytotoxicity.

Multi-specific antibody engineering is challenging because multiple domains must fold correctly without mutual interference. We designed an anti-ACE2 scFv via IASO, achieving exceptional structural confidence (pTM 0.9173, pLDDT 0.9157; **Supplementary Fig. 4b**), and linked it to an anti-CD3 scFv from Blinatumomab^87^ (**Fig. 4d**). Ternary complex prediction (BiTE-ACE2-CD3 epsilon) yielded high confidence scores (pLDDT 0.8612), indicating proper folding of both domains with preserved functional geometry for bispecific engagement (**Fig. 4e**).

Finally, we assessed developability using Blinatumomab as a clinical benchmark^87^. Aggregation analysis via Aggrescan4D^77^ showed that the IASO-designed BiTE maintains favorable Aggrescan4D score comparable to Blinatumomab (**Supplementary Fig. 4c**). Solubility predictions (CamSol method)^78^ exceeded Blinatumomab by 10% (**Supplementary Fig. 4d**), and immunogenicity assessment revealed reduced HLA-II binding for 10 out of 12 major HLA alleles compared to the approved therapeutic (**Supplementary Fig. 4e**). These lines of evidence demonstrate that IASO functions as a comprehensive platform for complex biopharmaceutical engineering, generating structurally and immunologically optimized multi-specific candidates that meet clinical development standards.

## Discussion

In this study, we developed IASO, an integrated computational platform that enables rational design of functional scFvs from antigen sequence alone, without requiring structural information. This framework addresses two fundamental challenges in therapeutic scFv discovery: the astronomical scale of explorable sequence space and the inefficiency of empirical screening. By integrating IASO-Gen for antigen-directed generation with IASO-AAI for high-precision interaction prediction, IASO transforms stochastic scFv discovery into predictable computational engineering. We demonstrated successful application across oncology targets, viral variants, and multi-specific formats, establishing IASO as a versatile platform for next-generation biopharmaceutical development.

A key innovation of IASO-AAI lies in its multi-scale integration of evolutionary and physicochemical information. While structure prediction models such as Boltz-2^75^ offer atomic-level accuracy, their computational demands prohibit exhaustive screening of large libraries. IASO-AAI addresses this limitation by combining protein language model embeddings, which capture global evolutionary patterns accumulated across millions of natural antibody sequences, with CKSAAP descriptors^44^ encoding local physicochemical rules governing hydrophobic and electrostatic interactions. This synergistic architecture enables rapid filtering that transcends simple sequence matching, achieving SOTA performance while maintaining computational efficiency essential for practical library screening. The dominance of antibody-specific features in our SHAP analysis reveals that specialized immunoglobulin grammar, rather than general protein features, drives binding prediction, highlighting the importance of domain-specific language models in biological sequence analysis.

The capacity of IASO-Gen to generate functional antibodies without structural data addresses a critical limitation of current computational design approaches. Structure-based methods (e.g., RFAntibody^30^) remain restricted to well-characterized targets with high-resolution structures as scaffolds, precluding application to intrinsically disordered regions, membrane proteins, or newly emerging viral variants where structural information is unavailable. By conditioning generation solely on antigen sequence via prefix-tuning and LoRA, IASO-Gen preserves the broad antibody grammar encoded in ProGen2-OAS^27^ while directing synthesis toward target-specific subspaces. Remarkably, IASO-designed candidates targeting the EGFR S468R mutant overcome steric clashes that abolish Cetuximab binding, demonstrating that sequence-based conditioning can achieve atomic-level structural optimization. This suggests that antigen sequence information, when properly integrated through prefix-tuning, implicitly encodes sufficient constraints to guide structurally compatible design.

The practical value of IASO emerges from synergistic integration of its components into a unified pipeline. While individual tools for generation or prediction exist, IASO’s hierarchical workflow—generation, interaction screening, and structural filtering—enables systematic enrichment of functional candidates while optimizing computational resources. This integration proved essential for addressing complex challenges including drug resistance and multi-specific engineering. The EGFR S468R case study demonstrates that the pipeline can rationally design resistance-overcoming scFvs by capturing subtle structural perturbations, while the BiTE application shows extension to complex formats requiring coordinated function of multiple domains. Critically, IASO-designed candidates achieved developability metrics comparable to or exceeding clinical benchmarks (Cetuximab-derived scFv, Blinatumomab), suggesting that the framework generates not merely binding sequences but pharmaceutically viable leads.

Despite these advancements, several limitations should be addressed. Current evaluations rely on *in silico* predictions and require experimental validation through biological assays such as ELISA or surface plasmon resonance^88^ to confirm binding kinetics and therapeutic efficacy in cellular models. Additionally, while our curated dataset mitigates bias, iterative refinement through Design-Build-Test-Learn cycles^89,90^ will be essential for enhancing model robustness. Integration of experimental feedback via active learning^91^ strategies should improve predictive and generative accuracy, particularly for underrepresented target classes. Future development should also address prediction of more complex properties including tissue penetration, pharmacokinetics, and in vivo stability. Expansion to additional antibody formats (e.g., nanobodies, IgGs) and incorporation of developability constraints directly into the generative objective could further streamline the path from computational design to clinical candidates.

In conclusion, IASO provides a rapid, rational platform for therapeutic antibody design from sequence data alone, addressing critical medical needs including drug-resistant mutants and pandemic viruses. We anticipate that IASO will become integral to AI-driven drug discovery, enabling efficient design of safe, efficacious biopharmaceuticals while substantially reducing development time and cost.

## Materials and Methods

### IASO-AAI model development

#### Dataset curation for IASO-AAI

We curated an antibody-antigen interaction (AAI) dataset from two public databases: AACDB^39^ (7,498 pairs) and AbRank^40^ (342,356 pairs). From the combined 349,854 pairs, we extracted VH and VL antibody sequences paired with their cognate antigen sequences. After removing entries with duplicate antigen sequences, we obtained 4,340 unique positive antibody–antigen pairs, corresponding to 4,340 unique antigens.

To generate biologically meaningful negative examples, we performed homology-aware negative sampling. Antigen sequences were clustered using CD-HIT^41^ at 90% sequence identity to group similar proteins. For each positive pair (antibody with antigen *X*), we generated five negative pairs by pairing the same antibody with randomly selected antigens *Y* from clusters different from *X*, following previous studies^42,43^. This strategy yields negative examples that are more biologically plausible than arbitrary mismatches, while reducing redundancy from highly similar antigen sequences. Consequently, it better reflects practical screening settings in which antibodies must be distinguished from multiple plausible non-cognate antigens. The final dataset comprised 26,040 pairs with a 1:5 positive-to-negative ratio, randomly partitioned into training (80%), validation (10%), and test (10%) sets.

#### Feature Engineering

IASO-AAI integrates three complementary feature representations capturing evolutionary context and local physicochemical properties. The first component, antibody embeddings, encodes concatenated VH and VL sequences using the antibody-specific language model AbLang-2^22^, yielding 480-dimensional vectors that capture immunoglobulin-specific evolutionary constraints and structural motifs. The second component, antigen embeddings, processes antigen sequences through ESM-2 (15B parameters)^21^ to generate 5,120-dimensional representations encoding general protein evolutionary patterns and functional constraints.

The third component, CKSAAP (Composition of k-Spaced Amino Acid Pairs) features^44^, complements these global evolutionary representations by quantifying local amino acid pair frequencies. For a sequence *s* = *s*_1_, *s*_2_, . . ., *s*_*L*_ of length L, the normalized frequency of an amino acid pair (*x*, *y*) separated by *k* residues is:

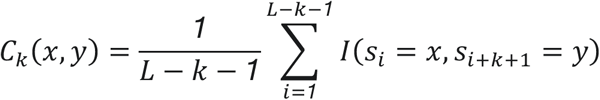

where *I* is the indicator function. We computed CKSAAP features for VH, VL, and antigen sequences with *k* ∈ {0, 1, 2, 3} following established protocols^23,92^. For each sequence, this produced a 1,600-dimensional vector (20^2^ amino acid pair types × 4 spacing values of *k*). Concatenating features from the VH sequence, VL sequence, and antigen sequence yielded 4,800 total CKSAAP dimensions (1,600 × 3). The final 10,400-dimensional feature vector concatenates all three representations (antibody embeddings: 480, antigen embeddings: 5,120, CKSAAP features: 4,800), enabling the model to leverage both macro-level evolutionary patterns and micro-level physicochemical interaction rules.

#### Training and performance evaluation of IASO-AAI

We trained a gradient-boosted decision tree model (LightGBM)^93^ on the integrated feature set. Hyperparameters were optimized using Bayesian optimization (Optuna)^94^to maximize the precision-recall area under the curve (PR-AUC) on the validation dataset. The optimized model was evaluated on a held-out test dataset, using PR-AUC, receiver operating characteristic AUC (ROC-AUC), accuracy, precision, recall, F1-score, and Matthews correlation coefficient (MCC).

For comparative benchmarking, we retrained five recent sequence-based AAI predictors (AbAgIntPre^46^, AntiBinder^48^, DeepInterAware^24^, MambaAAI^49^, and RLEAAI^23^) using their published implementations and identical train/validation/test splits as IASO-AAI.

To elucidate the contribution of different information sources, we performed feature importance analysis using SHAP (SHapley Additive exPlanations)^55^. For each feature type (antibody embeddings, antigen embeddings, and CKSAAP), we calculated mean absolute SHAP values by averaging across all constituent dimensions. Additionally, we conducted ablation studies by retraining seven different model variants: three single-feature models (antibody only, antigen only, CKSAAP only), three two-feature combinations, and the full model. All variants used hyperparameters optimized for the full model and were evaluated across three random seeds (42, 43, 44) to assess stability.

To verify generalization beyond our curated dataset, we evaluated IASO-AAI on an independent external dataset from RLEAAI^23^. We retrained IASO-AAI from scratch using the training dataset from the RLEAAI study^23^, following their exact data preprocessing and splitting protocols to ensure fair comparison. Performance on their held-out test set was compared against five established methods (AbAgIPA^43^, AbAgIntPre^46^, S3AI^53^, DeepAAI^54^, and RLEAAI^23^) previously benchmarked on this dataset, using PR-AUC, ROC-AUC, precision, recall, and F1-score as evaluation metrics.

#### Probability calibration and reliability evaluation

To assess whether the IASO-AAI outputs represent reliable probabilities rather than arbitrary confidence scores, we evaluated model calibration on the held-out test set. Predicted probabilities were binned into 10 equal intervals, and for each bin we compared the mean predicted probability (confidence) against the observed fraction of true positives, visualized through reliability diagrams. Calibration quality was quantified using two metrics: the Brier score^50^, which measures the mean squared error between predicted probabilities and binary outcomes (1 for binding, 0 for non-binding), and the expected calibration error (ECE)^51,52^, which quantifies the average absolute discrepancy between predicted confidence and empirical accuracy across bins.

### Antigen-conditioned scFv generation with IASO-Gen

#### Autoregressive foundation

IASO-Gen extends ProGen2-OAS^27^, obtained by fine-tuning the autoregressive protein language model ProGen2^27^ on antibody sequences from the OAS database^56^. The autoregressive architecture generates sequences by predicting each amino acid conditioned on preceding context, with generative probability factorizing as 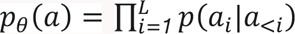 for sequence *a* = (*a*_1_, . . ., *a*_*L*_) . Training minimizes negative log-likelihood over a given protein dataset *D* = =*a^1^*, . . ., *a*^|*D*|^>:

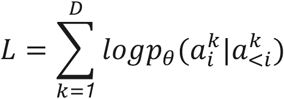

ProGen2 captures general protein evolutionary patterns, which are specialized to immunoglobulin grammar through OAS fine-tuning. However, ProGen2-OAS does not support generation conditioned on specific target antigens, which we address through antigen-conditional fine-tuning.

#### Training dataset curation

We constructed a paired antibody-antigen dataset from AACDB^39^ (7,498 pairs) and AbRank^40^ (342,356 pairs). From AACDB, we extracted VH and VL sequences identified by "heavy" and "light" descriptors, paired them using PDB identifiers, and merged with corresponding antigen sequences. From AbRank, we extracted VH, VL, and antigen sequence columns, filtered to retain only standard 20-amino-acid antigens, and removed duplicate VH-VL-antigen triplets. Combining both sources yielded 150,620 unique antibody-antigen pairs for training IASO-Gen.

#### Antigen-conditional fine-tuning

To enable target-specific generation, we fine-tuned ProGen2-OAS using parameter-efficient methods that preserve the base model’s antibody grammar while incorporating antigen information. We employed Prefix-Tuning^57,59^ to inject antigen representations as prefixes into attention layers, combined with Low-Rank Adaptation (LoRA)^59^ to enhance sequence expressivity with limited training data.

Antigen sequences were encoded via ESM-2 (15B)^21^ to produce embeddings, which were projected through a trainable multilayer perceptron into prefix vectors. These prefixes were prepended to the Key and Value matrices across all attention layers and heads, while Query matrices remained unchanged, enabling the model to consistently condition on antigen context during generation. Critically, the main ProGen2-OAS^27^ parameters remained frozen, and only the prefix projector and the LoRA adapters were trained, preserving pretrained antibody knowledge while learning antigen-specific binding preferences.

Unlike prior approaches^28^ that generate antigen sequences as part of the output, IASO-Gen conditions solely on antigen prefixes without reconstructing the antigen itself. This design circumvents context length constraints, allowing conditioning on antigens up to 1,024 residues (ESM-2’s maximum input length) rather than being limited by the model’s context window minus antibody length. The generation process factorizes as:

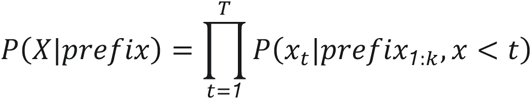

where *X* = (*x*_1_, . . ., *x*_*T*_) represents the generated antibody tokens and *prefix*_1:*k*_ is the antigen-derived prefix of length *k* Sequences were generated using the template <|*bos*|>-<*VH*>-[VH sequence]-<*SEP*>-<*VL*>-[VL sequence]-<|*eos*|>, where special tokens (<*VH*>, <*VL*>, <*SEP*>) were added to the ProGen2 tokenizer to delineate variable regions. Training optimized cross-entropy loss over antibody tokens only, excluding the antigen prefix from gradient computation.

#### scFv sequence generation

For inference, we combined the frozen ProGen2-OAS backbone with trained LoRA adapters and the antigen prefix projector. Target antigen sequences were encoded via ESM-2 and projected into prefixes for attention Key-Value matrices across all layers. Generation proceeded autoregressively from the template <*|bos|*>-<*VH*>-[VH sequence]-<*SEP*>-<*VL*>-[VL sequence]-<*|eos|*> using multinomial sampling with temperature scaling and nucleus (Top-p) sampling^95^.

To ensure physiologically realistic variable region lengths, we introduced a two-stage length constraint mechanism. When the VH chain reached a target length randomly sampled from *U*(110, 130) residues, we forcibly inserted <*SEP*><*VL*> tokens to initiate VL generation. Similarly, when VL reached a length sampled from *U*(105, 110), we inserted the <*|eos|*> to terminate generation, yielding sequences with VH: 110-130 aa and VL: 105-110 aa. These bounds were selected based on the canonical size of immunoglobulin variable domains and the empirically broader length variability of VH relative to VL, while remaining consistent with the IMGT V-domain numbering framework^96,97^.

#### Characterization of generated scFvs

We evaluated IASO-Gen by generating 10,000 antibody sequences conditioned on the EGFR antigen and comparing them against unconditioned baselines (ProGen2^27^, ProGen2-OAS^27^), the antigen-conditioned model (MAGE^28^), and 10,000 sequences randomly sampled from the OAS database^56^ as a reference for natural antibody properties.

To assess physicochemical characteristics, we computed molecular weight (MW), isoelectric point (pI), GRAVY score (Grand Average of Hydropathy, related to antibody aggregation)^60^, aromaticity, aliphatic index^61^, instability index, and net charge at pH 7.4 using Biopython^98^. We quantified similarity to natural antibodies through an OAS Reproducibility Score defined as 1-*D*_*ks*_, where *D*_*ks*_is the Kolmogorov–Smirnov statistic^99^ comparing each property’s distribution against the OAS reference.

Immunogenicity risk was assessed by predicting HLA class II binding affinity using the NetMHCIIpan 4.3 server^63^ for two representative HLA-DRB1 alleles (DRB1*04:01, DRB1*15:01)^64–66^. Each sequence was segmented into overlapping 15-residue peptides, and we quantified the number of strong binders (SB; rank score < 1%) per sequence.

The complementarity-determining region H3 (CDRH3), critical for antigen recognition^67,68^, was extracted using ANARCI with IMGT numbering^100^. We analyzed CDRH3 length distributions and assessed sequence diversity through Shannon entropy at each position^69,101^:

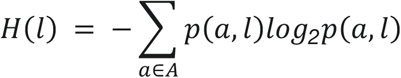

where *p*(*a*, *l*) is the probability of amino acid *a* at position *l* in the IMGT numbering, and through pairwise Levenshtein edit distances between CDRH3 sequences. Statistical comparisons employed Wilcoxon signed-rank tests for entropy distributions and Mann-Whitney U tests^102^ for edit distances.

To visualize the latent sequence space, we embedded generated antibodies using AbLang-2^22^ followed by UMAP projection^72^. To assess antigen-specific conditioning, we generated 10,000 sequences for each of four antigens (EGFR, PD-1, HIV gp120, SARS-CoV-2 spike) and evaluated whether IASO-Gen produces antigen-specific clustering in the projected sequence space.

### Design and validation of EGFR-targeting scFvs

To validate the IASO pipeline, we designed scFvs targeting human EGFR, a well-characterized therapeutic target in oncology^76,103^. We retrieved the EGFR sequence from PDB (4UV7) and generated the clinically relevant S468R resistant mutant^73^ by replacing serine 468 with arginine. As a benchmark, we constructed a Cetuximab-derived scFv by linking its heavy and light chains (retrieved from the Thera-SAbDab database^104^) with a standard (GGGGS)3 linker.

Using IASO-Gen, we encoded the EGFR antigen sequence via ESM-2 (15B), projected it into antigen prefixes, and generated 20,000 unique VH-VL pairs. These were linked with (GGGGS)3 to form scFv candidates, excluding sequences whose CDR regions could not be annotated by ANARCI^100^. For comparison, we generated equivalent libraries using MAGE^28^ (antigen-conditioned) and ProGen2^27^ (unconditioned). We screened all candidates using IASO-AAI, which outputs binding scores from 0 to 1, classifying sequences with scores >0.5 as predicted binders. Score distributions were compared across models to assess whether antigen conditioning enriched for EGFR-specific candidates.

To refine predictions and account for structural compatibility, we selected the top 100 scFv candidates (top 1% by IASO-AAI score) and predicted their complex structures with EGFR using Boltz-2^75^. Boltz-2 provides confidence metrics including pLDDT (local confidence), pTM (global fold accuracy), and pde (predicted distance error). We compared the distribution of these metrics between IASO-Gen and baseline candidates, and extracted sequences meeting stringent structural criteria (pTM > 0.7, pLDDT > 0.8) as high-confidence anti-EGFR scFvs. Selected complexes were visualized using PyMOL and compared against a Boltz-2 prediction of a previously reported anti-EGFR scFv^76^.

To assess whether IASO-designed scFvs possess pharmaceutical-grade properties, we performed *in silico* developability profiling using Boltz-2-predicted structures and compared candidates against the Cetuximab-derived scFv benchmark. Aggregation propensity was evaluated using Aggrescan4D^77^, which computes structurally corrected aggregation scores accounting for residue spatial arrangement. Average aggregation scores per residue were compared across candidates, with more negative values indicating lower aggregation risk and higher stability. Solubility was quantified using the CamSol Structurally Corrected tool^78^, which assigns profile scores where values >1 indicate high solubility and values <−1 indicate poor solubility.

Immunogenicity risk was assessed by predicting HLA class II binding affinity using NetMHCIIpan 4.3^63^ for 12 representative HLA-DRB1 alleles (DRB1*01:01, 03:01, 04:01, 07:01, 08:01, 09:01, 10:01, 11:01, 12:01, 13:01, 14:01, 15:01) covering major global populations^79–81^. ScFv sequences were segmented into overlapping 15-residue peptides, and binding affinity (rank score) was computed for each allele. We quantified potential T-cell epitopes by counting peptides classified as strong binders (rank < 1%) or weak binders (rank < 5%), comparing epitope burden against Cetuximab-derived scFv to assess relative immunogenicity risk.

### Versatility across diverse antigens and viral variants

To assess the generalizability of the IASO pipeline to structurally diverse targets, we designed antibodies against antigens spanning oncology (PD-1^82^, PDB: 3RRQ), infectious disease (HIV gp120^105^, PDB: 2NY3; SARS-CoV-2 variants), and host receptors (ACE2, PDB: 9ELE chain A). To evaluate variant discrimination capability, we targeted the receptor-binding domains (RBDs) of two closely related SARS-CoV-2 Omicron sublineages, JN.1 (PDB: 9IU1) and its derivative KP.3.1.1 (PDB: 9ELE, chain B). Although the two variants differ by only two residues overall, our structural analysis focused on the substitution at position 175 (Gln→Glu), which represented the key local difference discussed here. Antigen sequences were retrieved from PDB, purification tags removed, and encoded via ESM-2 (15B) to generate antigen prefixes for IASO-Gen.

For each antigen, we generated scFv libraries using IASO-Gen, screened candidates via IASO-AAI binding score predictions, and predicted complex structures of top-scoring candidates using Boltz-2^75^. To assess variant specificity, we performed cross-binding analysis for scFvs designed against the KP.3.1.1 RBD by predicting their complex structures with both KP.3.1.1 (intended target) and JN.1 (parental strain) using Boltz-2. Interface interactions were analyzed using the Molecular Operating Environment (MOE) software, focusing on the local environment surrounding residue 175 to identify binding mode differences, hydrogen bonding patterns, and steric clashes that discriminate between Glu175 (KP.3.1.1) and Gln175 (JN.1).

### Expansion to bispecific T-cell engagers

To demonstrate the extensibility of the IASO pipeline to multi-specific formats, we designed a bispecific T-cell engager (BiTE)^7–9^ comprising an IASO-generated anti-ACE2 scFv (target recognition domain) linked via a (GGGGS)3 linker to an anti-CD3 scFv sequence derived from Blinatumomab^87^ (T-cell recruitment domain). We predicted the ternary complex structure (BiTE–ACE2–CD3 epsilon) using Boltz-2 by simultaneously inputting the BiTE sequence, ACE2 (PDB: 9ELE chain A), and CD3 epsilon (PDB: 6JXR chain D). Structural integrity of the BiTE polypeptide was assessed via pLDDT scores across the entire molecule.

Developability and immunogenicity of the designed BiTE were evaluated using the same protocols applied to single scFvs, with Blinatumomab serving as the clinical benchmark. The Boltz-2-predicted BiTE structure was analyzed for aggregation propensity (Aggrescan4D^77^), solubility (CamSol^78^), and immunogenicity risk (NetMHCIIpan 4.3^63^ across 12 HLA-DRB1 alleles), quantifying strong and weak HLA-II binders as potential T-cell epitopes.

### Statistical analysis

Statistical analyses were performed using SciPy (v1.15.2) in Python (v3.10.18). Differences in physicochemical properties of generated antibodies were assessed using two-sided Kolmogorov–Smirnov tests for all pairwise comparisons, with Holm correction for multiple testing. For IASO-AAI score comparisons, homogeneity of variance was first assessed via Levene’s test, followed by one-way ANOVA (equal variance) or Welch’s one-way ANOVA (unequal variances). Distributions of Boltz-2 structural metrics were compared using Cramér–von Mises and Epps–Singleton tests for overall distributional differences, Mann–Whitney U tests for location shifts (with Cliff’s delta effect sizes), and Levene’s test for variance differences. Multiple comparisons were corrected using the Holm (physicochemical properties) or Benjamini-Hochberg FDR (Boltz-2 metrics) procedures. Statistical significance was defined as α = 0.05 for unadjusted tests and adjusted p < 0.05 or q < 0.05 for corrected tests.

### Computational environment

All computational procedures in this study were performed on the TSUBAME 4.0 supercomputer at the Institute of Science Tokyo. Each compute node comprises two Intel Xeon CPU Max 9480 processors (112 cores total) and four NVIDIA H100 SXM5 GPUs.

## ACKNOWLEDGEMENTS

This work was supported by KAKENHI grants from the Japan Society for the Promotion of Science (JSPS) to H.S. (23K28184, 24H01755 and 25H01571), the JST FOREST Program to H.S. (JPMJFR242Q), as well as a grant from the Canon Foundation. We thank Y. Otani, T. Sakuma, T. Suzuki, H. Hishinuma, and T. Ito for critical reading of the manuscript, K. Tanaka for help with preparation of the manuscript, and all laboratory members for helpful discussions. Figures were created in part with BioRender.com.

## AUTHOR CONTRIBUTIONS

H.S. designed and supervised the study. T.F. developed the framework, performed all formal analyses, and drafted the manuscript. H.S. and T.F. revised and finalized the manuscript. All authors have read and approved the final manuscript.

## References

1. Crescioli, S. et al. Antibodies to watch in 2026. MAbs 18(1) 2614669, (2026).

2. Chan, A. C., Martyn, G. D. & Carter, P. J. Fifty years of monoclonals: the past, present and future of antibody therapeutics. Nature Reviews Immunology vol. 25 745–765 (2025).

3. Lu, R. M. et al. Technological advancements in antibody-based therapeutics for treatment of diseases. J. Biomed. Sci. 32, 98 (2025).

4. Ahmad, Z. A. et al. scFv Antibody: Principles and Clinical Application. J. Immunol. Res. 2012, 980250 (2012).

5. Zhang, F. et al. Single chain fragment variable (scFv) antibodies targeting the spike protein of porcine epidemic diarrhea virus provide protection against viral infection in piglets. Viruses 11, (2019).

6. Muñoz-López, P. et al. Single-Chain Fragment Variable: Recent Progress in Cancer Diagnosis and Therapy. Cancers 14(17), 4206 (2022).

7. Huehls, A. M., Coupet, T. A. & Sentman, C. L. Bispecific T-cell engagers for cancer immunotherapy. Immunol. Cell Biol. 93, 290–296 (2015).

8. van de Donk, N. W. C. J. & Zweegman, S. T-cell-engaging bispecific antibodies in cancer. The Lancet 402, 142–158 (2023).

9. Brinkmann, U. & Kontermann, R. E. The making of multispecific immunoglobulins–a clinical perspective. mAbs 18(1), 2613548 (2026).

10. Eshhar, Z., Waks, T., Gross, G. & Schindler, D. G. Specific activation and targeting of cytotoxic lymphocytes through chimeric single chains consisting of antibody-binding domains and the gamma or zeta subunits of the immunoglobulin and T-cell receptors. Proceedings of the National Academy of Sciences 90, 720–724 (1993).

11. Sterner, R. C. & Sterner, R. M. CAR-T cell therapy: current limitations and potential strategies. Blood Cancer J. 11, 69 (2021).

12. Klein, C., Brinkmann, U., Reichert, J. M. & Kontermann, R. E. The present and future of bispecific antibodies for cancer therapy. Nature Reviews Drug Discovery 23(4), 301–319 (2024).

13. Pansri, P., Jaruseranee, N., Rangnoi, K., Kristensen, P. & Yamabhai, M. A compact phage display human scFv library for selection of antibodies to a wide variety of antigens. BMC Biotechnol. 9, 6 (2009).

14. Zambrano, N. et al. High-Throughput Monoclonal Antibody Discovery from Phage Libraries: Challenging the Current Preclinical Pipeline to Keep the Pace with the Increasing mAb Demand. Cancers vol. 14 Preprint at 10.3390/cancers14051325 (2022).

15. Han, K. H., Li, Y.-C., Parveen, R., Venkataraman, S. & Lin, C.-W. Technologies for Monoclonal Antibody Discovery and Development. Int. J. Mol. Sci. 26, 10470 (2025).

16. Khan, A. et al. Toward real-world automated antibody design with combinatorial Bayesian optimization. Cell Reports Methods 3(1), 100374 (2023).

17. Norman, R. A. et al. Computational approaches to therapeutic antibody design: Established methods and emerging trends. Brief. Bioinform. 21, 1549–1567 (2020).

18. Cheng, J., Liang, T., Xie, X. Q., Feng, Z. & Meng, L. A new era of antibody discovery: an in-depth review of AI-driven approaches. Drug Discov. Today 29(6), 103984 (2024).

19. Kim, D. N., McNaughton, A. D. & Kumar, N. Leveraging Artificial Intelligence to Expedite Antibody Design and Enhance Antibody–Antigen Interactions. Bioengineering 11, 185 (2024).

20. Graves, J. et al. A Review of Deep Learning Methods for Antibodies. Antibodies 2020, Vol. 9, Page 12 9, 12 (2020).

21. Lin, Z., et al. Evolutionary-Scale Prediction of Atomic-Level Protein Structure with a Language Model. Science vol. 379 https://www.science.org (2023).

22. Olsen, T. H., Moal, I. H. & Deane, C. M. Addressing the antibody germline bias and its effect on language models for improved antibody design. Bioinformatics 40, (2024).

23. Hu, J., Zhou, Y., Zhang, W. Y. & Zhou, X. G. RLEAAI: improving antibody-antigen interaction prediction using protein language model and sequence order information. Brief. Bioinform. 26(3), bbaf238 (2025).

24. Xia, Y. et al. DeepInterAware: Deep Interaction Interface-Aware Network for Improving Antigen-Antibody Interaction Prediction from Sequence Data. Advanced Science 12(13), e2412533 (2025).

25. Vaswani, A. et al. Attention Is All You Need. In Advances in Neural Information Processing Systems 30 (NeurIPS 2017) (2017).

26. Hummer, A. M., Schneider, C., Chinery, L. & Deane, C. M. Investigating the volume and diversity of data needed for generalizable antibody–antigen ΔΔG prediction. Nat. Comput. Sci. 5, 635–647 (2025).

27. Nijkamp, E., Ruffolo, J. A., Weinstein, E. N., Naik, N. & Madani, A. ProGen2: Exploring the boundaries of protein language models. Cell Syst. 14, 968–978.e3 (2023).

28. Wasdin, P. T. et al. Generation of antigen-specific paired-chain antibodies using large language models. Cell 188(25), 7206–7221.e16 (2025).

29. Stark, H. et al. BoltzGen: Toward Universal Binder Design. bioRxiv 2025.11.20.689494 (2025). 10.1101/2025.11.20.689494.

30. Bennett, N. R. et al. Atomically accurate *de novo* design of antibodies with RFdiffusion. Nature 649, 183–193 (2026).

31. Watson, J. L., et al. *De novo* design of protein structure and function with RFdiffusion. Nature 620, 1089–1100 (2023).

32. Zheng, S. J. Deep learning for antigen-specific antibody design. Veterinary Molecular Immunology 425–442 (2026) doi:10.1007/978-981-99-8929-4_5.

33. Meng, F. et al. A comprehensive overview of recent advances in generative models for antibodies. Computational and Structural Biotechnology Journal vol. 23 2648–2660 Preprint at 10.1016/j.csbj.2024.06.016 (2024).

34. Zheng, J., Wang, Y., Liang, Q., Cui, L. & Wang, L. The Application of Machine Learning on Antibody Discovery and Optimization. Molecules 29, 5923 (2024).

35. Kong, Y. et al. A synergistic generative-ranking framework for tailored design of therapeutic single-domain antibodies. Cell Discov. 11, 85 (2025).

36. Svilenov, H. L., Arosio, P., Menzen, T., Tessier, P. & Sormanni, P. Approaches to expand the conventional toolbox for discovery and selection of antibodies with drug-like physicochemical properties. mAbs 15(1), 2164459 (2023).

37. Matsunaga, R. & Tsumoto, K. Accelerating antibody discovery and optimization with high-throughput experimentation and machine learning. Journal of Biomedical Science 32(1), 46 (2025).

38. Santuari, L., Bachmann Salvy, M., Xenarios, I. & Arpat, B. AI-accelerated therapeutic antibody development: practical insights. Frontiers in Drug Discovery 4, (2024).

39. Zhou, Y. et al. AACDB: Antigen-Antibody Complex Database — a Comprehensive Database Unlocking Insights into Interaction Interface. eLife reviewed preprint 104934 (2025).

40. Liu, C., et al. AbRank: A Benchmark Dataset and Metric-Learning Framework for Antibody-Antigen Affinity Ranking. arXiv 2506.17857 (2025).

41. Li, W. & Godzik, A. Cd-hit: A fast program for clustering and comparing large sets of protein or nucleotide sequences. Bioinformatics 22, 1658–1659 (2006).

42. Sun, C. et al. SAGERank: inductive learning of protein–protein interaction from antibody–antigen recognition. Chem. Sci. 16, 17885–17899 (2025).

43. Gu, M., Yang, W. & Liu, M. Prediction of antibody-antigen interaction based on backbone aware with invariant point attention. BMC Bioinformatics 25, (2024).

44. Chen, K., Kurgan, L. A. & Ruan, J. Prediction of flexible/rigid regions from protein sequences using k-spaced amino acid pairs. BMC Struct. Biol. 7, (2007).

45. Wang, X.-B., Wu, L.-Y., Wang, Y.-C. & Deng, N.-Y. Prediction of palmitoylation sites using the composition of k-spaced amino acid pairs. Protein Engineering Design and Selection 22, 707–712 (2009).

46. Huang, Y., Zhang, Z. & Zhou, Y. AbAgIntPre: A deep learning method for predicting antibody-antigen interactions based on sequence information. Front. Immunol. 13, (2022).

47. Zhou, H.-X. & Pang, X. Electrostatic Interactions in Protein Structure, Folding, Binding, and Condensation. Chem. Rev. 118, 1691–1741 (2018).

48. Zhang, K., Tao, Y. & Wang, F. AntiBinder: utilizing bidirectional attention and hybrid encoding for precise antibody–antigen interaction prediction. Brief. Bioinform. 26(1), bbaf008 (2025).

49. Liu, X., Fu, H., Yang, Y. & Zhang, J. Bio-Inspired Mamba for Antibody–Antigen Interaction Prediction. Biomolecules 15(6), 764 (2025).

50. Glenn, W. B. Verification of forecasts expressed in terms of probability. Mon. Weather Rev. 78, 1–3 (1950).

51. Naeini, M. P., Cooper, G. F. & Hauskrecht, M. Obtaining Well Calibrated Probabilities Using Bayesian Binning. In Proceedings of the AAAI Conference on Artificial Intelligence 29(1), 2901–2907 (2015).

52. Guo, C., Pleiss, G., Sun, Y. & Weinberger, K. Q. On Calibration of Modern Neural Networks. http://arxiv.org/abs/1706.04599 (2017).

53. Liu, Y. et al. Interpretable antibody-antigen interaction prediction by introducing route and priors guidance. Preprint at 10.1101/2024.03.09.584264 (2024).

54. Zhang, J. et al. Predicting unseen antibodies’ neutralizability via adaptive graph neural networks. *Nat*. Mach. Intell. 4, 964–976 (2022).

55. Lundberg, S. M. & Lee, S.-I. A Unified Approach to Interpreting Model Predictions. In Advances in Neural Information Processing Systems 30 (2017).

56. Olsen, T. H., Boyles, F. & Deane, C. M. Observed Antibody Space: A diverse database of cleaned, annotated, and translated unpaired and paired antibody sequences. Protein Science 31, 141–146 (2022).

57. Li, X. L. & Liang, P. Prefix-Tuning: Optimizing Continuous Prompts for Generation. In Proceedings of the 59th Annual Meeting of the Association for Computational Linguistics and the 11th International Joint Conference on Natural Language Processing (Volume 1: Long Papers), 4582–4597 (2021).

58. Luo, J. et al. Controllable Protein Design by Prefix-Tuning Protein Language Models. Preprint at 10.1101/2023.12.03.569747 (2023).

59. Hu, E. J., et al. LoRA: Low-Rank Adaptation of Large Language Models. In ICLR 2022 (2022).

60. Kyte, J. & Doolittle, R. F. A simple method for displaying the hydropathic character of a protein. J. Mol. Biol. 157, 105–132 (1982).

61. Ikai, A. Thermostability and aliphatic index of globular proteins. J. Biochem. 88, 1895–8 (1980).

62. Harris, C. T. & Cohen, S. Reducing Immunogenicity by Design: Approaches to Minimize Immunogenicity of Monoclonal Antibodies. BioDrugs vol. 38 205–226 Preprint at 10.1007/s40259-023-00641-2 (2024).

63. Nilsson, J. B. et al. Accurate prediction of HLA class II antigen presentation across all loci using tailored data acquisition and refined machine learning. Sci. Adv. 9(47), eadj6367 (2023).

64. Dessen, A., Lawrence, C. M., Cupo, S., Zaller, D. M. & Wiley, D. C. X-Ray Crystal Structure of HLA-DR4 (DRA*0101, DRB1*0401) Complexed with a Peptide from Human Collagen II. Immunity 7, 473–481 (1997).

65. Anderson, K. M. et al. A Molecular Analysis of the Shared Epitope Hypothesis: Binding of Arthritogenic Peptides to DRB1*04 Alleles. Arthritis & Rheumatology 68, 1627–1636 (2016).

66. Alcina, A. et al. Multiple Sclerosis Risk Variant HLA-DRB1*1501 Associates with High Expression of DRB1 Gene in Different Human Populations. PLoS One 7, e29819 (2012).

67. Harding, F. A., Stickler, M. M., Razo, J. & DuBridge, R. B. The immunogenicity of humanized and fully human antibodies: Residual immunogenicity resides in the CDR regions. MAbs 2, 256–265 (2010).

68. Wu, T. Te, Johnson, G. & Kabat, E. A. Length distribution of CDRH3 in antibodies. Proteins: Structure, Function, and Bioinformatics 16, 1–7 (1993).

69. Shannon, C. E. A Mathematical Theory of Communication. Bell System Technical Journal 27, 379–423 (1948).

70. Wagner, R. A. & Fischer, M. J. The String-to-String Correction Problem. Journal of the ACM 21, 168–173 (1974).

71. Waury, K., Lelieveld, S., Abeln, S. & van den Ham, H.-J. Comparison of sequence- and structure-based antibody clustering approaches on simulated repertoire sequencing data. PLoS Comput. Biol. 21, e1013057 (2025).

72. McInnes, L., Healy, J., Saul, N. & Großberger, L. UMAP: Uniform Manifold Approximation and Projection. J. Open Source Softw. 3, 861 (2018).

73. García-Foncillas, J. et al. Distinguishing Features of Cetuximab and Panitumumab in Colorectal Cancer and Other Solid Tumors. Front. Oncol. 9, (2019).

74. Li, S. et al. Structural basis for inhibition of the epidermal growth factor receptor by cetuximab. Cancer Cell 7, 301–311 (2005).

75. Passaro, S., et al. Boltz-2: Towards Accurate and Efficient Binding Affinity Prediction. bioRxiv 2025.06.14.659707 (2025). doi:10.1101/2025.06.14.659707.

76. Liu, X. et al. A cross-reactive pH-dependent EGFR antibody with improved tumor selectivity and penetration obtained by structure-guided engineering. Mol. Ther. Oncolytics 27, 256–269 (2022).

77. Bárcenas, O. et al. Aggrescan4D: structure-informed analysis of pH-dependent protein aggregation. Nucleic Acids Res. 52, W170–W175 (2024).

78. Sormanni, P., Aprile, F. A. & Vendruscolo, M. The CamSol Method of Rational Design of Protein Mutants with Enhanced Solubility. J. Mol. Biol. 427, 478–490 (2015).

79. Gregersen, P. K. HLA class II polymorphism: implications for genetic susceptibility to autoimmune disease. Lab. Invest. 61, 5–19 (1989).

80. Tiercy, J.-M. Molecular basis of HLA polymorphism: implications in clinical transplantation. Transpl. Immunol. 9, 173–180 (2002).

81. Lannan, M. B. et al. A universal monoallelic human leukocyte antigen class II immunopeptidomic platform for defining the immunogenicity potential of therapeutic proteins. Commun. Biol. 8, 1243 (2025).

82. Ishida, Y. PD-1: Its Discovery, Involvement in Cancer Immunotherapy, and Beyond. Cells 9, 1376 (2020).

83. Liu, Y. et al. Lineage-specific pathogenicity, immune evasion, and virological features of SARS-CoV-2 BA.2.86/JN.1 and EG.5.1/HK.3. Nat. Commun. 15, 8728 (2024).

84. Shi, J. et al. Pathogenicity, virological features, and immune evasion of SARS-CoV-2 JN.1-derived variants including JN.1.7, KP.2, KP.3, and KP.3.1.1. Nat. Commun. 16, 11002 (2025).

85. Damas, J. et al. Broad host range of SARS-CoV-2 predicted by comparative and structural analysis of ACE2 in vertebrates. Proceedings of the National Academy of Sciences 117, 22311–22322 (2020).

86. Monteil, V. et al. Clinical grade ACE2 as a universal agent to block SARS-CoV-2 variants. EMBO Mol. Med. 14, (2022).

87. Przepiorka, D. et al. FDA Approval: Blinatumomab. Clinical Cancer Research 21, 4035–4039 (2015).

88. Homola, J. Surface Plasmon Resonance Sensors for Detection of Chemical and Biological Species. Chem. Rev. 108, 462–493 (2008).

89. Gill, R. T., Halweg-Edwards, A. L., Clauset, A. & Way, S. F. Synthesis aided design: The biological design-build-test engineering paradigm? Biotechnol. Bioeng. 113, 7–10 (2016).

90. Carbonell, P. et al. An automated Design-Build-Test-Learn pipeline for enhanced microbial production of fine chemicals. *Commun*. Biol. 1, 66 (2018).

91. Tharwat, A. & Schenck, W. A Survey on Active Learning: State-of-the-Art, Practical Challenges and Research Directions. Mathematics 11, 820 (2023).

92. Ye, C., Hu, W. & Gaeta, B. Prediction of Antibody-Antigen Binding via Machine Learning: Development of Data Sets and Evaluation of Methods. JMIR Bioinform. Biotech. 3, (2022).

93. Ke, G. et al. LightGBM: A Highly Efficient Gradient Boosting Decision Tree. https://github.com/Microsoft/LightGBM.

94. Akiba, T., Sano, S., Yanase, T., Ohta, T. & Koyama, M. Optuna: A Next-generation Hyperparameter Optimization Framework. http://arxiv.org/abs/1907.10902 (2019).

95. Holtzman, A., Buys, J., Du, L., Forbes, M. & Choi, Y. The Curious Case of Neural Text Degeneration. (2020).

96. Chiu, M. L., Goulet, D. R., Teplyakov, A. & Gilliland, G. L. Antibody Structure and Function: The Basis for Engineering Therapeutics. Antibodies (Basel) 8, (2019).

97. Lefranc, M.-P. et al. IMGT unique numbering for immunoglobulin and T cell receptor variable domains and Ig superfamily V-like domains. Dev. Comp. Immunol. 27, 55–77 (2003).

98. Cock, P. J. A. et al. Biopython: Freely available Python tools for computational molecular biology and bioinformatics. Bioinformatics 25, 1422–1423 (2009).

99. Massey, F. J. The Kolmogorov-Smirnov Test for Goodness of Fit. J. Am. Stat. Assoc. 46, 68 (1951).

100. Dunbar, J. & Deane, C. M. ANARCI: antigen receptor numbering and receptor classification. Bioinformatics 32, 298–300 (2016).

101. Shenkin, P. S., Erman, B. & Mastrandrea, L. D. Information-theoretical entropy as a measure of sequence variability. Proteins: Structure, Function, and Bioinformatics 11, 297–313 (1991).

102. Mann, H. B. & Whitney, D. R. On a Test of Whether one of Two Random Variables is Stochastically Larger than the Other. The Annals of Mathematical Statistics 18, 50–60 (1947).

103. Banisadr, A., Safdari, Y., Kianmehr, A. & Pourafshar, M. Production of a germline-humanized cetuximab scFv and evaluation of its activity in recognizing EGFR-overexpressing cancer cells. Hum. Vaccin. Immunother. 14, 856–863 (2018).

104. Raybould, M. I. J. et al. Thera-SAbDab: the Therapeutic Structural Antibody Database. Nucleic Acids Res. 48, D383–D388 (2020).

105. Pantophlet, R. & Burton, D. R. GP120: Target for Neutralizing HIV-1 Antibodies. Annu. Rev. Immunol. 24, 739–769 (2006).

