## Supplementary Tables S1-S3, Supplementary Figure S1-S4 for "*De novo* design of therapeutic scFvs and multi-specific engagers from sequence alone"

**Table 1 | Hyperparameter optimization for IASO-AAI**

| Parameter | Search Space (Range) | Optimized Value |
|---|---|---|
| learning_rate | LogUniform (0.001, 0.2) | 0.0187 |
| num_leaves | IntLogUniform (10, 1024) | 57 |
| max_depth | IntUniform (3, 12) | 9 |
| min_data_in_leaf | IntUniform (10, 300) | 23 |
| feature_fraction | Uniform (0.6, 1.0) | 0.621 |
| bagging_fraction | Uniform (0.6, 1.0) | 0.985 |
| bagging_freq | IntUniform (0, 10) | 7 |
| lambda_l1 | LogUniform (1.0×10-8, 10) | $2.32 \times 10^{-6}$ |
| lambda_l2 | LogUniform (1.0×10-8, 10) | $4.75 \times 10^{-6}$ |

**Table 2 | Performance comparison of AAI prediction models**

| Model | PR-AUC | ROC-AUC | Accuracy | Precision | Recall | F1 | MCC |
|---|---|---|---|---|---|---|---|
| IASO-AAI (**ours**) | **0.7495** | **0.9311** | **0.8829** | **0.8525** | 0.3594 | 0.5057 | **0.5059** |
| AbAgIntPre[46] | 0.3052 | 0.7449 | 0.5472 | 0.2611 | **0.9378** | 0.4084 | 0.3094 |
| AntiBinder[48] | 0.4324 | 0.7775 | 0.8294 | 0.4716 | 0.3410 | 0.3958 | 0.3048 |
| DeepInterAware[24] | 0.6008 | 0.8688 | 0.8580 | 0.5727 | 0.5820 | **0.5773** | 0.4920 |
| MambaAAI[49] | 0.5076 | 0.8305 | 0.8455 | 0.5166 | 0.4660 | 0.4900 | 0.4000 |
| RLEAAI[23] | 0.5692 | 0.8611 | 0.8652 | 0.6387 | 0.4944 | 0.5574 | 0.4849 |

**Table 3 | External validation of IASO-AAI on independent external dataset**

| Model | PR-AUC | ROC-AUC | Precision | Recall | F1 |
|---|---|---|---|---|---|
| IASO-AAI (**ours**) | **0.974** | **0.969** | **0.938** | 0.877 | **0.907** |
| AbAgIPA[43] | 0.781 | 0.721 | 0.810 | 0.555 | 0.654 |
| AbAgIntPre[46] | 0.739 | 0.694 | 0.666 | 0.619 | 0.640 |
| S3AI[53] | 0.940 | 0.938 | 0.853 | 0.863 | 0.857 |
| DeepAAI[54] | 0.946 | 0.945 | 0.842 | 0.903 | 0.872 |
| RLEAAI[23] | 0.939 | 0.948 | 0.887 | **0.911** | 0.899 |

### Supplementary Figure Legends

#### Supplementary Figure 1 | Technical validation of IASO-AAI

**(a)** Dataset construction pipeline. Antibody-antigen interaction (AAI) pairs from AACDB and AbRank databases were filtered and clustered via CD-HIT (90% identity threshold). Negative examples were generated through homology-aware sampling, yielding 26,040 pairs (1:5 positive:negative ratio). **(b)** CKSAAP *k*-mer parameter sensitivity. PR-AUC (left) and ROC-AUC (right) for $k$ = 1-5. Error bars: standard deviation ($n$=3 independent seeds). Performance remains stable across $k$ values; $k$ = 3 was selected following established computational benchmarks. **(c)** Confusion matrix on the independent test dataset ($n$=2,604 pairs, 1:5 positive:negative ratio, threshold 0.5). Rows: true labels; columns: predicted labels. The model achieves 98.9% specificity and 86.5% precision (154/178 predicted positives). **(d)** Probability calibration. Predicted probabilities binned into 10 intervals. x-axis: mean predicted probability, y-axis: observed positive fraction. Diagonal: perfect calibration. Low Brier score (0.079) and expected calibration error (ECE: 0.049) indicate reliable probability estimates. Histogram shows distribution of predicted probabilities.

#### Supplementary Figure 2 | Sequence diversity and latent space analysis of IASO-Gen

**(a)** Physicochemical properties for 10,000 antibodies from OAS database (reference) and four generative models (ProGen2, ProGen2-OAS, MAGE, and our IASO-Gen). Violin plots compare molecular weight (MW), aliphatic index, and net charge. **(b)** Shannon entropy across CDRH3 positions. IASO-Gen (blue) shows significantly higher sequence diversity than MAGE (red) ($p$ = 0.0422). **(c)** CDRH3 Levenshtein distance distributions. IASO-Gen (blue, mean 10.01) exhibits larger edit distances than MAGE (red, mean 9.09), indicating broader sequence space exploration (*** $p < 0.001$). **(d)** UMAP projection of antibody sequences embeddings targeting EGFR. IASO-Gen (blue) occupies a unique cluster distinct from ProGen2 (green), ProGen2-OAS (orange), and MAGE (red), suggesting access to unique antibody sequence space.

#### Supplementary Figure 3 | Interaction analysis and developability assessment of EGFR S468R-targeting scFv

**(a)** Interaction energetics at Arg468. Contact residues and calculated interaction energies for Cetuximab-derived scFv (left) versus IASO-designed scFv (right). Blue bars: favorable (negative) energies; red bars: unfavorable (positive) energies. **(b)** Aggregation propensity (Aggrescan4D)[77]. Structure-corrected aggregation scores for Cetuximab-derived scFv (–0.5559) and IASO scFv (average –0.4691). More negative scores indicate lower aggregation risk. The IASO scFv maintains a favorable aggregation-resistant profile comparable to the clinically approved therapeutic. **(c)** Solubility prediction (CamSol)[78]. Structure-corrected solubility scores; values >1 indicate high solubility. The IASO-designed scFv shows comparable solubility to the Cetuximab-derived scFv. **(d)** Immunogenicity assessment. Predicted HLA-II strong and weak binders across 12 HLA-DRB1 alleles. The IASO scFv (blue) exhibits binding frequencies equal to or lower than the Cetuximab-derived scFv (orange), indicating minimal immunogenicity risk.

#### Supplementary Figure 4 | Multi-antigen versatility and BiTE developability

**(a)** Structural metrics for IASO-designed scFvs targeting diverse antigens (PD-1, HIV gp120, SARS-CoV-2 KP.3.1.1 RBD, ACE2). Boltz-2-predicted complexes show high pLDDT (left), high pTM (middle), and low pde (right) across all targets, demonstrating framework versatility. **(b)** Anti-ACE2 scFv structure (BiTE target-recognition domain). High confidence scores (pTM 0.9173, pLDDT 0.9157) ensure stable multi-specific assembly. **(c)** Aggregation propensity (Aggrescan4D). IASO BiTE (lower, average –0.6509) shows comparable aggregation resistance to Blinatumomab (upper, average –0. 6970). More negative scores indicate lower aggregation risk. **(d)** Solubility (CamSol). The IASO BiTE (2.101) exhibits higher solubility than Blinatumomab (1.916); values >1 indicate high solubility. **(e)** Immunogenicity assessment. HLA-II binding predictions across 12 HLA-DRB1 alleles. The IASO BiTE (blue) shows binding frequencies at or below Blinatumomab (orange) for 10 out of 12 evaluated alleles, indicating minimal immunogenicity risk.

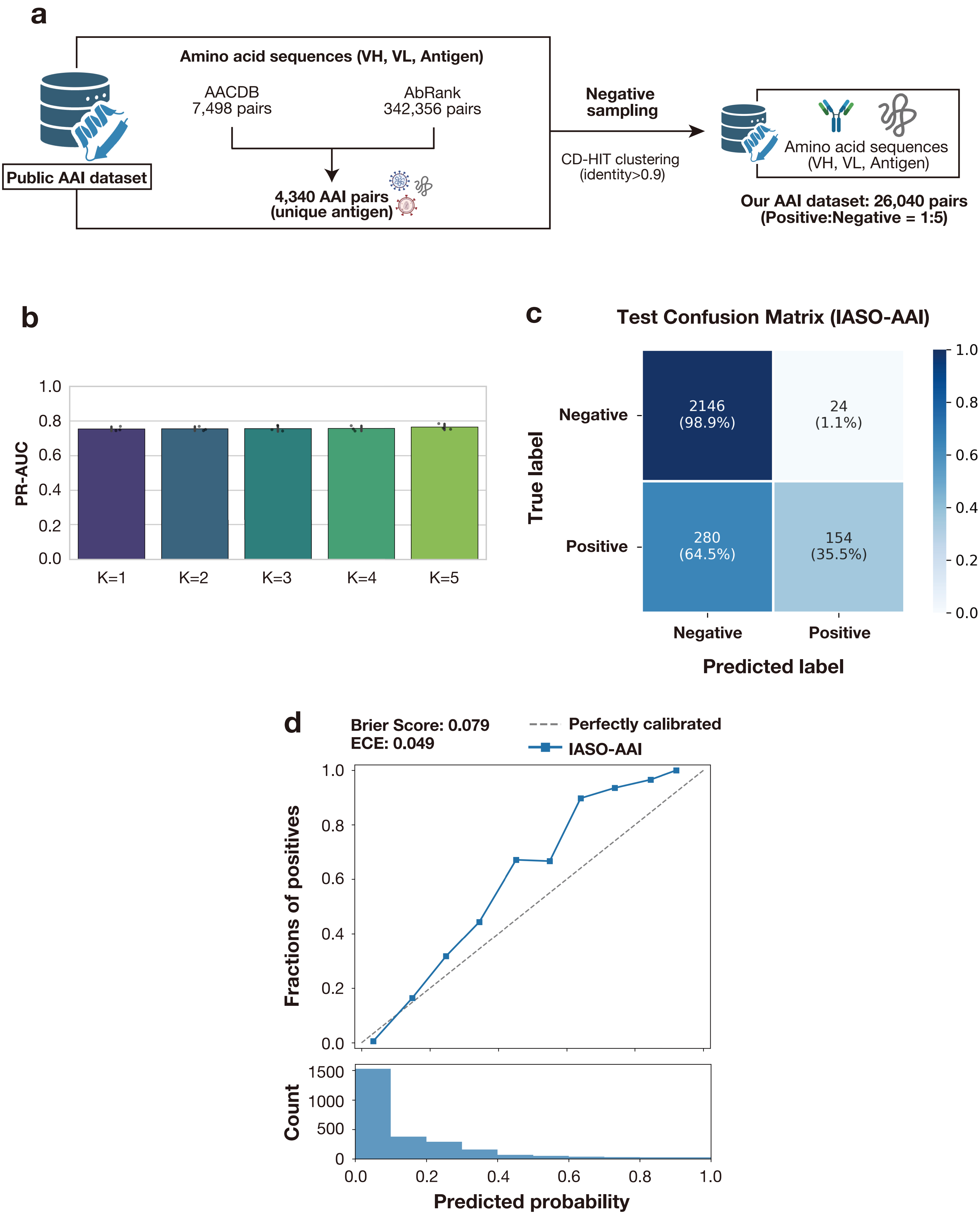


Fujiwara et al. Supplementary Figure 1

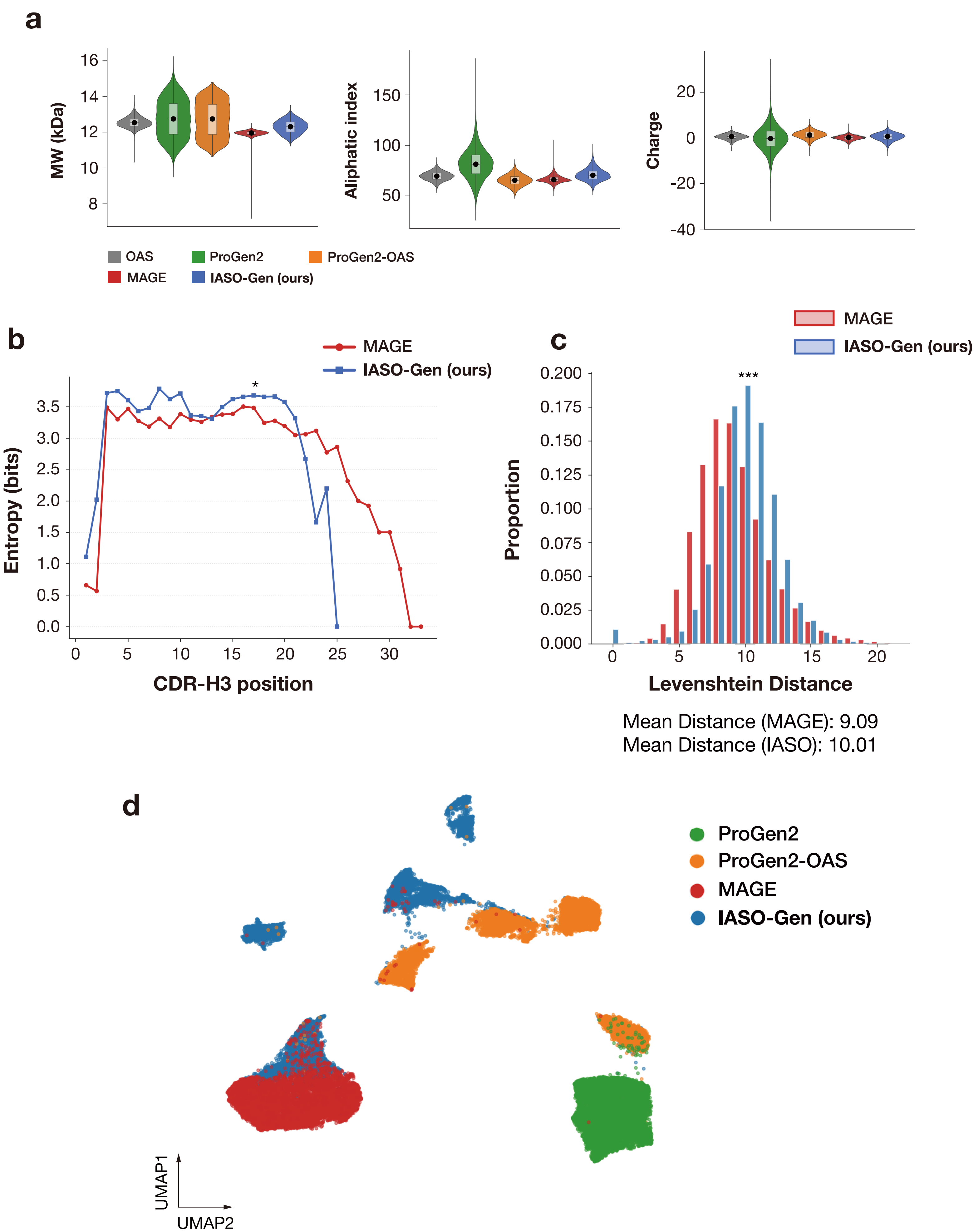


Fujiwara et al. Supplementary Figure 2

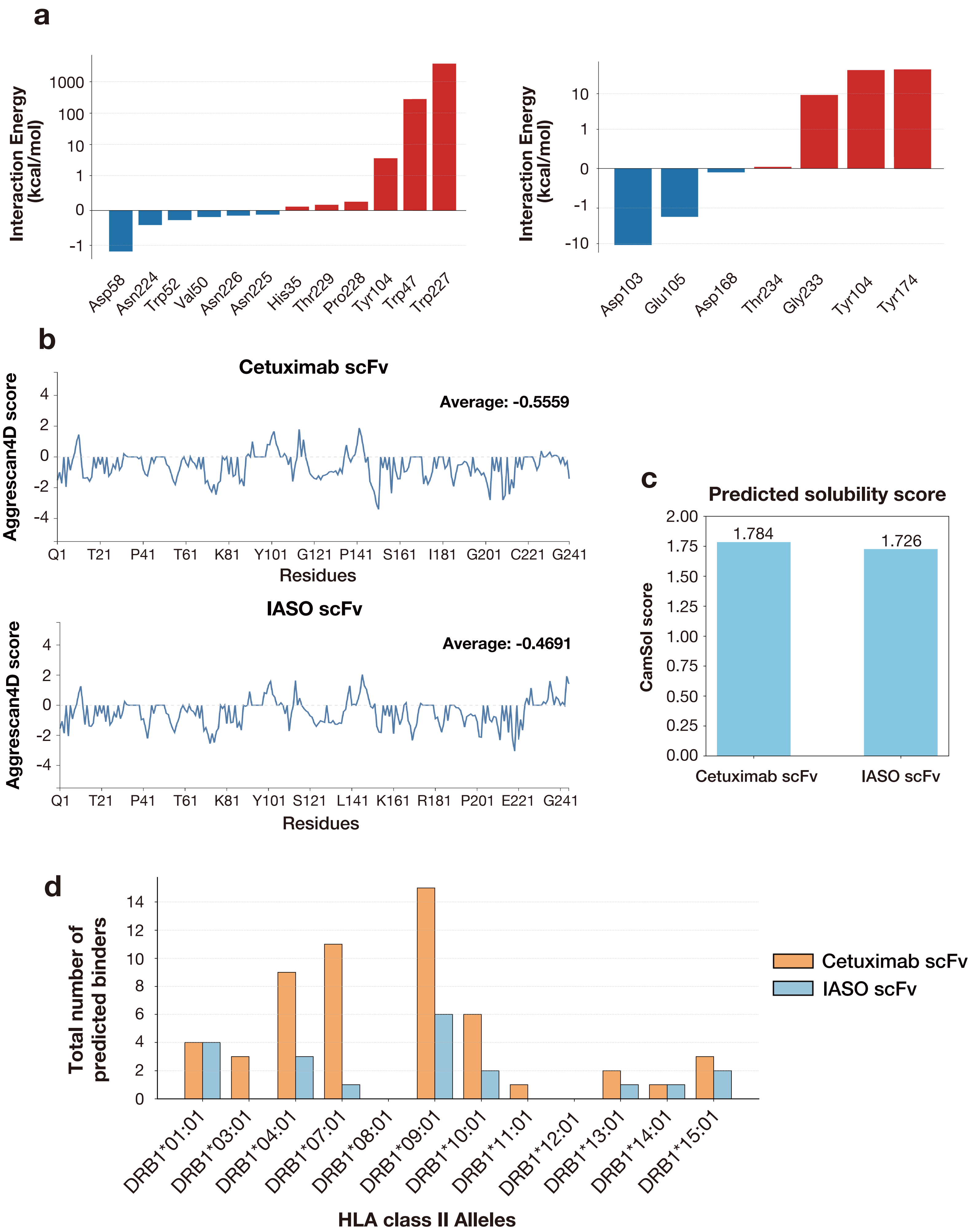


Fujiwara et al. Supplementary Figure 3

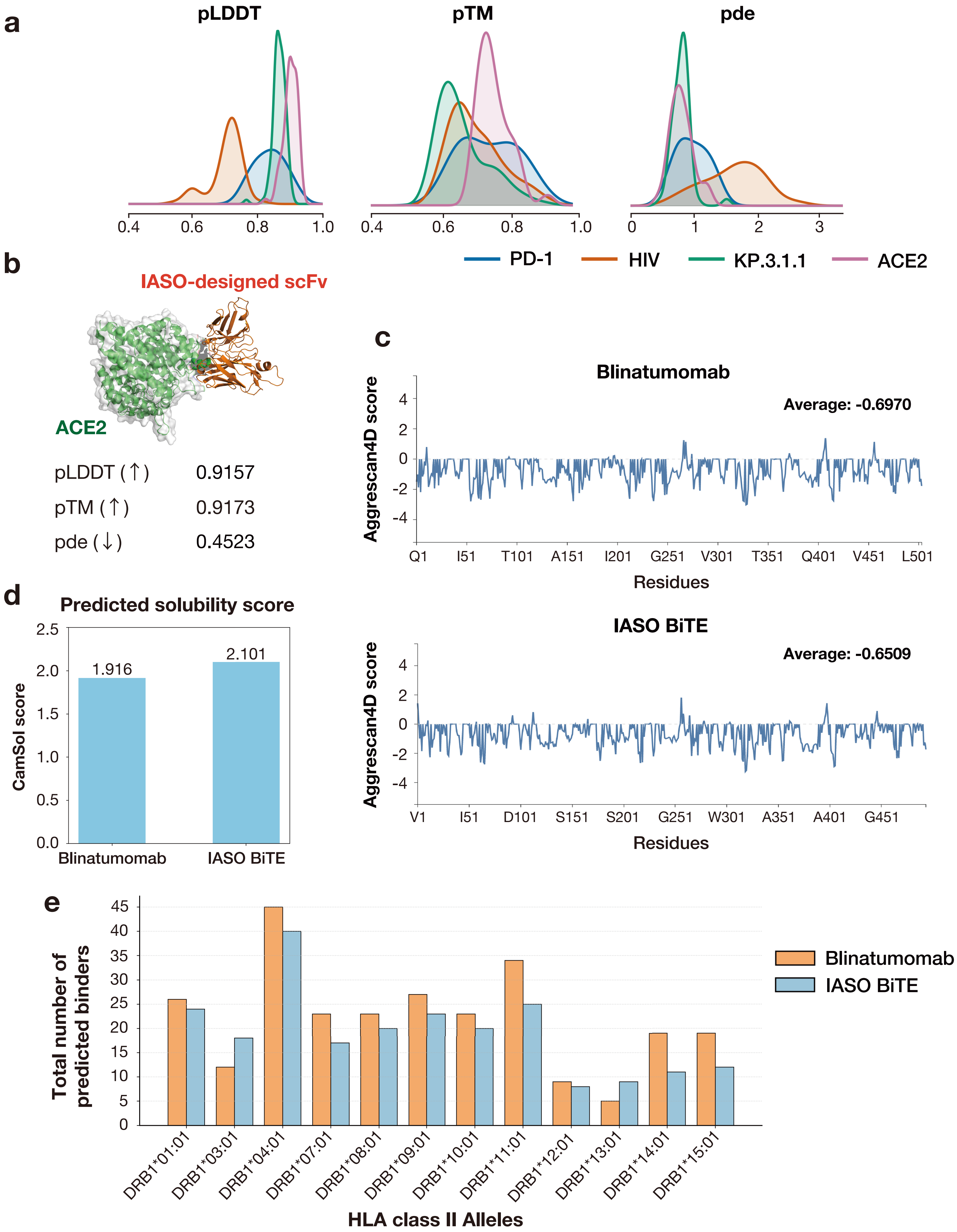


Fujiwara et al. Supplementary Figure 4
